# Quantitative modelling predicts the impact of DNA methylation on RNA polymerase II traffic

**DOI:** 10.1101/391904

**Authors:** Justyna Cholewa-Waclaw, Ruth Shah, Shaun Webb, Kashyap Chhatbar, Bernard Ramsahoye, Oliver Pusch, Miao Yu, Philip Greulich, Bartlomiej Waclaw, Adrian Bird

## Abstract

Patterns of gene expression are primarily determined by proteins that locally enhance or repress transcription. While many transcription factors target a restricted number of genes, others appear to modulate transcription levels globally. An example is MeCP2, an abundant methylated-DNA binding protein that is mutated in the neurological disorder Rett Syndrome. Despite much research, the molecular mechanism by which MeCP2 regulates gene expression is not fully resolved. Here we integrate quantitative, multi-dimensional experimental analysis and mathematical modelling to show that MeCP2 is a novel type of global transcriptional regulator whose binding to DNA creates "slow sites" in gene bodies. Waves of slowed-down RNA polymerase II formed behind these sites travel backward and indirectly affect initiation, reminiscent of defect-induced shock waves in non-equilibrium physics transport models. This mechanism differs from conventional gene regulation mechanisms, which often involve direct modulation of transcription initiation. Our findings uncover a genome-wide function of DNA methylation that may account for the reversibility of Rett syndrome in mice. Moreover, our combined theoretical and experimental approach provides a general method for understanding how global gene expression patterns are choreographed.

---

Many eukaryotic chromatin-associated factors modulate transcription by binding to specific sites in gene promoters or enhancers (1, 2). Most transcription factors are thought to modulate the initiation rate of transcription by altering histone-DNA interactions (2, 3) or imposing promoter-proximal obstacles (4). However, transcription can also be affected by processes that occur in the bodies of genes. In particular, DNA methylation, which is widespread in gene bodies, appears to affect progression of RNA polymerase II (RNA Pol II) through densely methylated exons (5). The mechanism is unclear, but methyl-CpG binding proteins (6) may be involved. Since most gene bodies contain methylated CpGs, such proteins may have a global effect on transcription.

One putative global modulator is methyl-CpG binding protein 2 (MeCP2) (7, 8), which is highly expressed in neurons. *MECP2* mutations, including loss-of-function or gene duplication, lead to severe neurological disorders (9, 10). MeCP2 does not behave as a conventional transcription factor with discrete targets, as its binding site occurs on average every ~100 base pairs. Evidence from *in vitro* (11, 12) and mouse models (13, 14) suggests that MeCP2 can mediate DNA methylation-dependent transcriptional inhibition. Transcriptional changes in mouse brain when MeCP2 is absent or over-expressed are relatively subtle but widespread (15–17), and the molecular mechanisms underlying these changes are unknown.

Here we set out to resolve the mechanism of MeCP2-dependent transcriptional regulation. Because MeCP2 binding sites occur in the vast majority of genes, we reasoned that most are likely to be influenced to some extent by its presence. To confront the technical and analytical challenges posed by modest changes in expression of large numbers of genes, we adopted a quantitative approach that combined deep, high quality datasets obtained from a uniform population of Lund Human Mesencephalic (LUHMES)-derived human dopaminergic neurons (18) with computational modelling. We created a spectrum of LUHMES cell lines expressing distinct levels of MeCP2. Using transposase-accessible chromatin sequencing (ATAC-seq) and chromatin immunoprecipitation (ChIP-seq) together with mathematical modelling, we detected a robust footprint of MeCP2 binding to mCG and mCA *in vivo* and determined the amount of MeCP2 bound to DNA. Quantification of mRNA abundance by RNA-seq revealed a relationship between changes in transcription and the density of mCG on gene bodies. To explain this observation, we proposed and tested several distinct mechanistic models. The only model consistent with our experimental results is one in which MeCP2 leads to slowing down of RNA polymerase II progression through a transcription unit. Importantly, mutant MeCP2 that is unable to bind NCoR fails to repress efficiently, suggesting that repression depends upon this interaction.

## Results

### Global changes in transcription correlate with MeCP2 expression level

We created progenitor cell lines capable of differentiation to a uniform population of human neurons (Fig. S1A-C) that expressed seven widely different levels of MeCP2, including knock-out (KO), wild-type (WT) and 11-fold over-expression (11x) (Figs. 1A,B and S1D; see Table S1). All lines differentiated into neurons with similar kinetics, expressed neuronal markers (Fig. S1E), and had identical global levels of DNA methylation (~3.7% of all cytosines were methylated) (Fig. S2A). To determine effects of MeCP2 levels on transcription, we performed RNA-seq on all seven cell lines. Most of the ∼ 17000 expressed genes responded to MeCP2 abundance, but changes were small (Fig. S3A). Based on known affinity of MeCP2 for methylated CG (mCG), we expected MeCP2 to affect genes depending on their mCG content. We therefore quantified DNA methylation for all genes in WT neurons using whole-genome bisulfite sequencing (TAB-seq) (Fig. S2B,C). In order to detect small changes in expression which might otherwise be obscured by other regulatory mechanisms and statistical noise, expressed genes were binned according to methylation density, considering gene bodies and promoters separately.

**Fig. 1.**
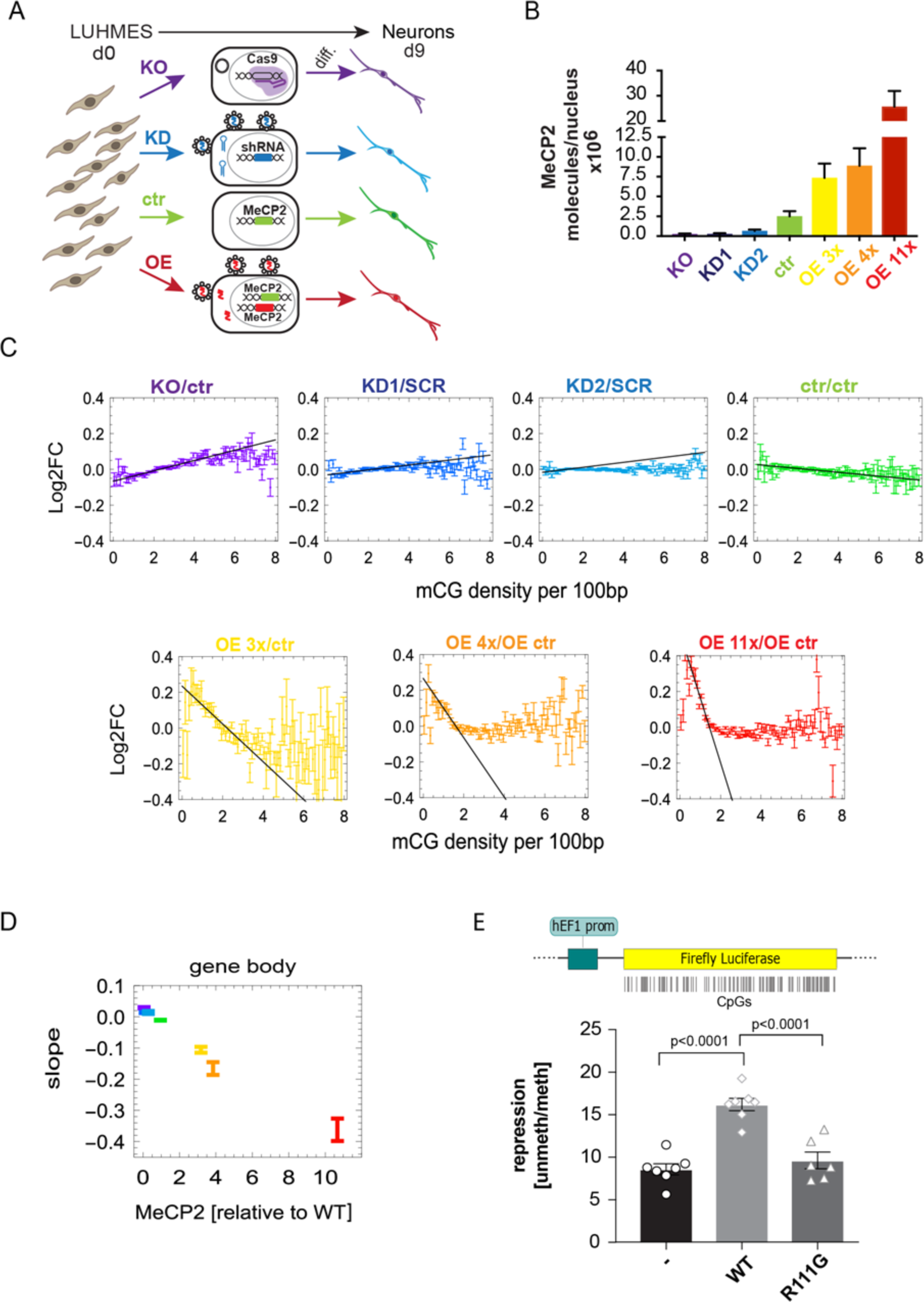
Gene expression strongly correlates with gene body mCG density and MeCP2 abundance. (A) Experimental design (Methods). (B) Mean number of MeCP2 molecules per nucleus. (C) Log2 fold change of gene expression (Log2FC) relative to appropriate controls (ctr – unmodified controls; SCR – scrambled shRNA control, OE ctr – overexpression control) for all seven levels of MeCP2, plotted against gene body mCG density. All Log2FC values have been shifted so that Log2FC averaged over all genes is zero. Black line indicates the maximum slope. (D) The maximum slope for gene bodies varies proportionally to MeCP2 abundance. (E) Ratio between luciferase expressions from an unmethylated and gene-body methylated constructs, for three cases: no MeCP2, WT MeCP2, and an MBD mutant R111G that is unable to bind mCG. Points show individual replicates. In all panels, error bars represent +/− SEM.

The average change in expression versus appropriate controls (Log2FC) showed a strong relationship to mCG density (*ρ*_mCG_) in gene bodies (Fig. 1C). The effect was the strongest for *ρ*_mCG_=0.8−4.0 mCG per 100bp which includes the vast majority of genes (Fig. S3B). Moreover, the maximum slope of the Log2FC versus *ρ*_mCG_ in gene bodies (Fig. 1C, black lines) was strikingly proportional to MeCP2 levels (Fig. 1D). In contrast, plots of Log2FC versus *ρ*_mCG_ in promoter regions showed a slope close to zero, indicating minimal dependence on promoter methylation (Fig. S3C). No clear dependence on MeCP2 level was observed for Log2FC versus total gene body mCG or mCG mean (Figs. S3D, E). These results indicated that the gene-body mCG density is the strongest predictor of MeCP2-dependent transcriptional changes. To test for a causal relationship, we transfected cells with two versions (methylated or unmethylated gene body) of a luciferase reporter gene with a methylation-free promoter in the presence of wildtype or the DNA binding mutant MeCP2[R111G] (Fig. S4). We observed a two-fold repression of methylated versus unmethylated luciferase gene body in the presence of WT MeCP2 compared to either no MeCP2 or mutant MeCP2 (Figs. 1E).

### MeCP2 binds predominantly methylated CG genome-wide

To map the binding of MeCP2 in human neurons, we performed MeCP2 ChIP-seq for KO, WT, OE 4x and OE 11x, and simultaneously developed a computer model that simulates the ChIP-seq procedure and MeCP2 binding *in vivo* (Fig. 2A). As expected, ChIP enrichment was proportional to the level of MeCP2 in each cell line (Fig. S5A-C) and showed a strong peak centred at mCGs in MeCP2-positive lines (Fig. 2B) as well as a correlation between MeCP2 enrichment and mCG density (Fig. 2C). Conversely, enrichment was absent at non-methylated CpG islands, non-methylated CGs, and GT dinucleotides (Fig. S5E,F).

**Fig 2.**
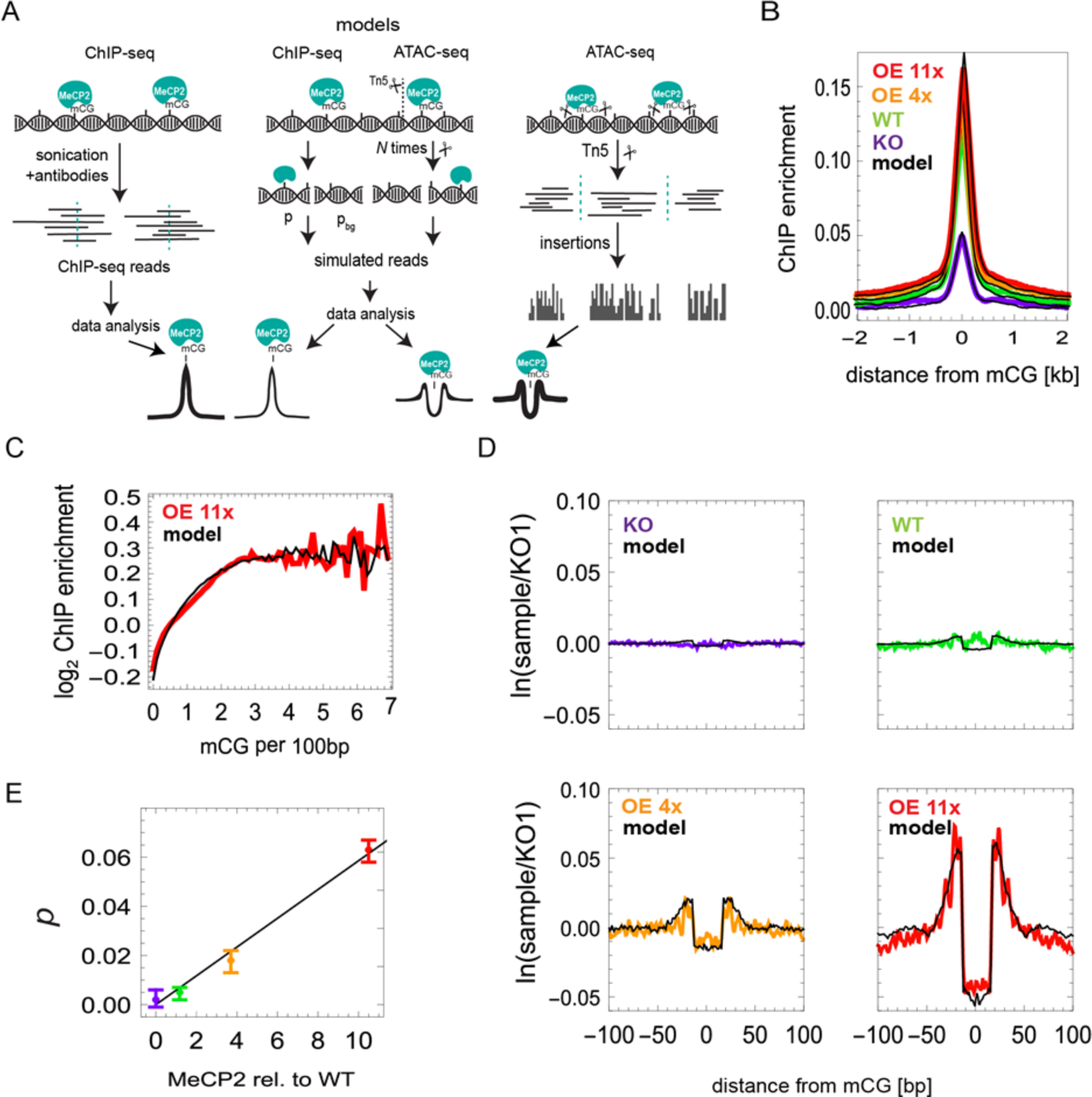
MeCP2 occupancy on the DNA is proportional to mCG density and MeCP2 level. (A) MeCP2 ChIP- and ATAC-seq experimental procedures and their *in silico* counterparts. *p*_*bg*_ and *p* are probabilities of background and MeCP2-bound reads, respectively. Tn5 insertion sites (scissors) occur in exposed DNA regions. (B) ChIP-seq enrichment profiles centred at mCG dinucleotides for different cell lines. Black lines represent *in silico* profiles fitted to the experimental data. (C) MeCP2 ChIP-seq enrichment data in OE 11x/KO (red) as a function of mCG density. (D) Average depletion profiles (logarithm of the ratio between the number of Tn5 insertions in a given cell line and KO1, 2-4 biological replicates) in the +/−100 bp regions surrounding mCG dinucleotides. Black lines represent computer simulations of the model fitted to the data. (E) Predicted fraction of mCGs occupied by MeCP2 versus MeCP2 level obtained from depletion profiles in (D). Error bars represent +/− SEM.

To derive an independent measure of absolute MeCP2 density on the DNA and to detect its molecular footprint with high resolution, we performed ATAC-seq (19) in which transposase Tn5 cuts exposed DNA to reveal DNA accessibility within chromatin (Fig. 2A). In agreement with ChIP-seq, ATAC-seq Tn5 insertion profiles (Figs. 2D) showed a graded depletion of insertion sites centered around mCG in WT, OE 4x and OE 11x neurons, whose amplitude was proportional to MeCP2 concentration (Fig. 2E) and therefore represents a “molecular footprint” of MeCP2 binding *in vivo.* An equivalent but weaker profile was seen at mCA, but was absent at non-methylated CG and GT and TA dinucleotides (Fig. S6). The size and amplitude of the footprint agrees well with a computer model of ATAC-seq and MeCP2 binding (Fig. 2D, black lines) and previous *in vitro* data (20, 21), confirming that MeCP2 occupies 11bp of DNA in living cells. The model revealed that only 6.3% of mCG sites are actually occupied by MeCP2 in *OE 11x* neurons, falling to less than 1% occupancy in *WT* (Fig. 2E), perhaps due in part to occlusion by nucleosomes. Excellent agreement between the models and ATAC-seq and ChIP-seq data allows us to predict MeCP2 occupancy from mCG density and MeCP2 level in each cell line.

### MeCP2 does not regulate transcription via condensation of chromatin or premature termination

To interpret these results mechanistically, we considered mathematical models based on a commonly accepted paradigm for gene expression (Fig. S7A) (22). In the first class of models named Condensation models (Fig. 3A), MeCP2 affects the rate of transcription initiation via changes in chromatin structure. The possibility that MeCP2 affects the initiation rate *α* by binding to promoters was rejected because it would imply a stronger correlation between gene expression and *ρ*_mCG_ in promoters than in gene bodies, contrary to our observations (Fig. S3C). MeCP2 could hypothetically affect the fraction *f* of cells with specific genes in the ON state via some long-distance mechanism involving binding to gene bodies and changing the degree of chromatin openness near promoters. However, mapping chromatin accessibility using ATAC-seq showed that while there is a correlation between MeCP2 and accessibility (Fig. 3B), it cannot account for the observed Log2FC in gene expression (Fig. 3C).

**Fig. 3.**
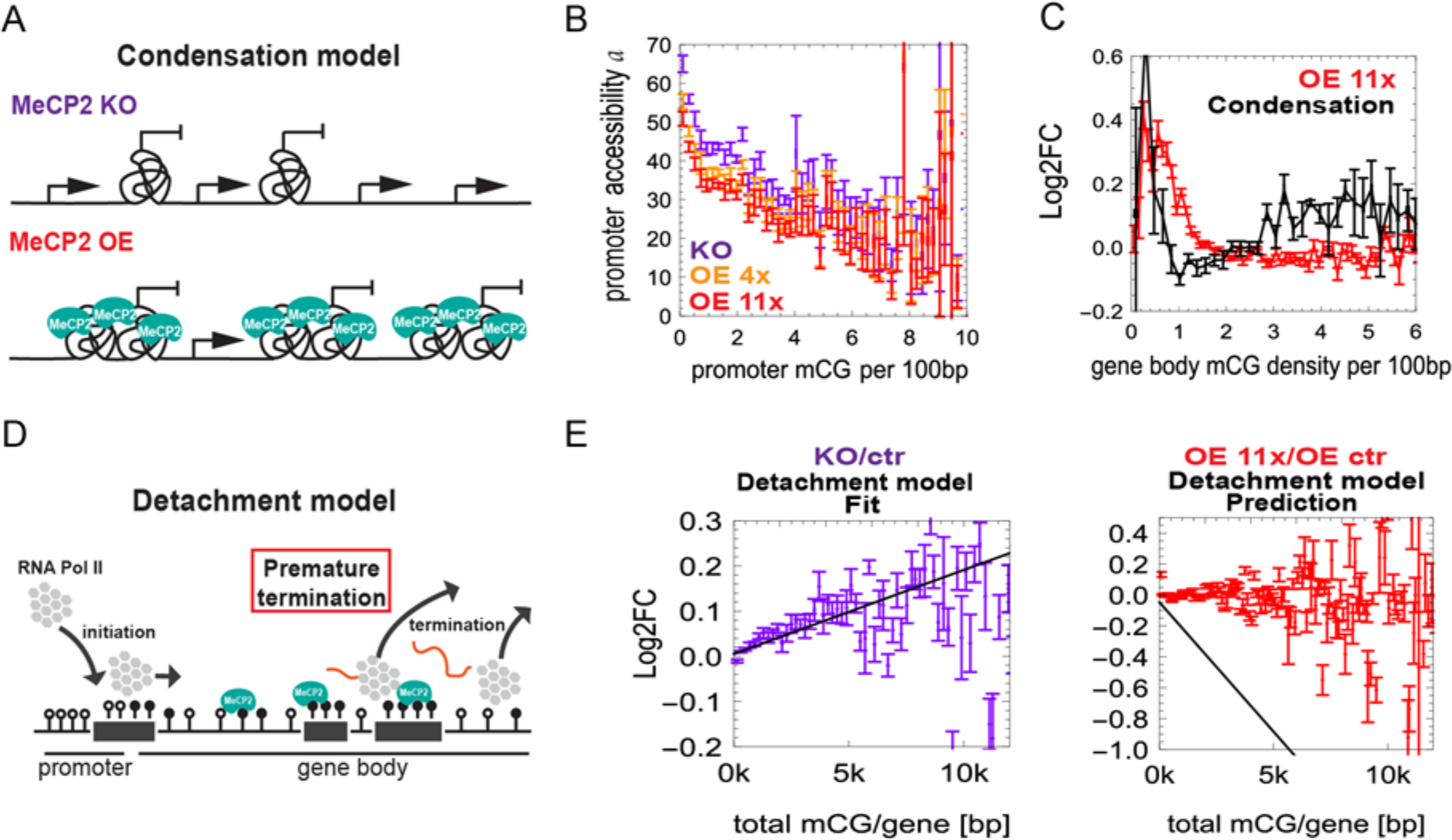
MeCP2 does not regulate transcription via condensation of chromatin or premature termination. (A) A cartoon of the Condensation model. Tangles represent regions of condensed chromatin that are inaccessible to RNA Pol II. (B) Chromatin accessibility (measured by ATAC-seq) at promoters decreases with promoter methylation but changes very little with MeCP2 level. (C) The Condensation model disagrees with Log2FC(OE 11x/KO) obtained from RNA-seq. (D) Schematic representation of the Detachment model. (E) Log2FC (gene expression) for: KO/ctr (left, purple) and 11x OE/OE ctr (right, red) versus the total number of mCGs in the gene. Black lines represent predictions of Detachment model. Error bars represent +/− SEM.

We next considered potential effects of MeCP2 on the elongation phase of transcription. The Detachment Model posits that MeCP2 causes transcription to prematurely abort (Fig. 3D). Since the probability of termination increases with each blocking site, under this model the Log2FC should increase in proportion to the number of methylated CGs (*N*_mCG_) in the gene and MeCP2 level, and the slope of Log2FC should be linear with respect to the MeCP2 level. These predictions are in disagreement with the data (Figs. 3E and S3D). Therefore, it is unlikely that MeCP2 affects transcription via premature termination.

### MeCP2 creates “Dynamical obstacles” that impede transcriptional elongation

Finally, we considered a “Congestion model” whereby Pol II pauses when it encounters MeCP2 itself or an induced, transient structural modification of chromatin (Fig. 4A). The parameters are: the fraction *p* of mCGs bound by MeCP2, MeCP2 turn-over (unbinding) rate *k*_*u*_, and (specific to each gene) the length *L* of the gene, the density *ρ*_mCG_ of methylated CGs, and the initiation rate *α*. Fig. 4B shows the transcription rate for OE 11x predicted by the model as a function of *α*, for different *ρ*_mCG_. The assumed value of *k*_*u*_ = 0.04 s^−1^is compatible with the reported *in vivo* residence time of MeCP2 on chromatin (25-40s (23)). Inspired by non-equilibrium statistical mechanics of one-dimensional transport models (24, 25), we expect a non-equilibrium phase transition from a low-density to a maximal-current (congested) phase as the initiation rate or the density of obstacles increase beyond a certain critical point. Indeed, all curves in Fig. 4B have a characteristic shape: a linear relationship *J* ≈ *α* for small *α*, followed by saturation at high initiation rates. Saturation occurs due to congestion as polymerases queue upstream of obstacles (SI Videos 1,2). However, even in the non-saturated regime of intermediate *α*, excluded-volume interactions between polymerases that have been slowed down by an obstacle cause a density shock wave that propagates backwards (Fig. 4C). A small increase in the density of polymerases near the promoter decreases the rate of Pol II binding to the TSS. Thus, even though MeCP2 does not directly affect Pol II initiation, it does so indirectly by shock waves that form behind MeCP2-induced obstacles in gene bodies (Fig. 4D). To test the model against RNA-seq data, we estimated average initiation rates for genes with similar mCG densities by fitting the model to Log2FC(OE 11x/OE ctr) versus *ρ*_mCG_ (Fig. 4E left, see also SI Section 10.2). We then used the model to predict Log2FC data for the remaining 6 cell lines. The model strikingly reproduces the data (Fig. 4E for OE 4x and KO) as well as the slopes of the Log2FC plots for all seven cell lines (Fig. 4F). A similar behaviour occurs in a modified model in which Pol II slows down (rather than completely stops) on permanent or long-lasting structural modifications of chromatin (Fig. S7B-E, SI Video 3). We conclude that both congestion models are compatible with the experimental data presented in Fig. 1C and D.

**Fig. 4.**
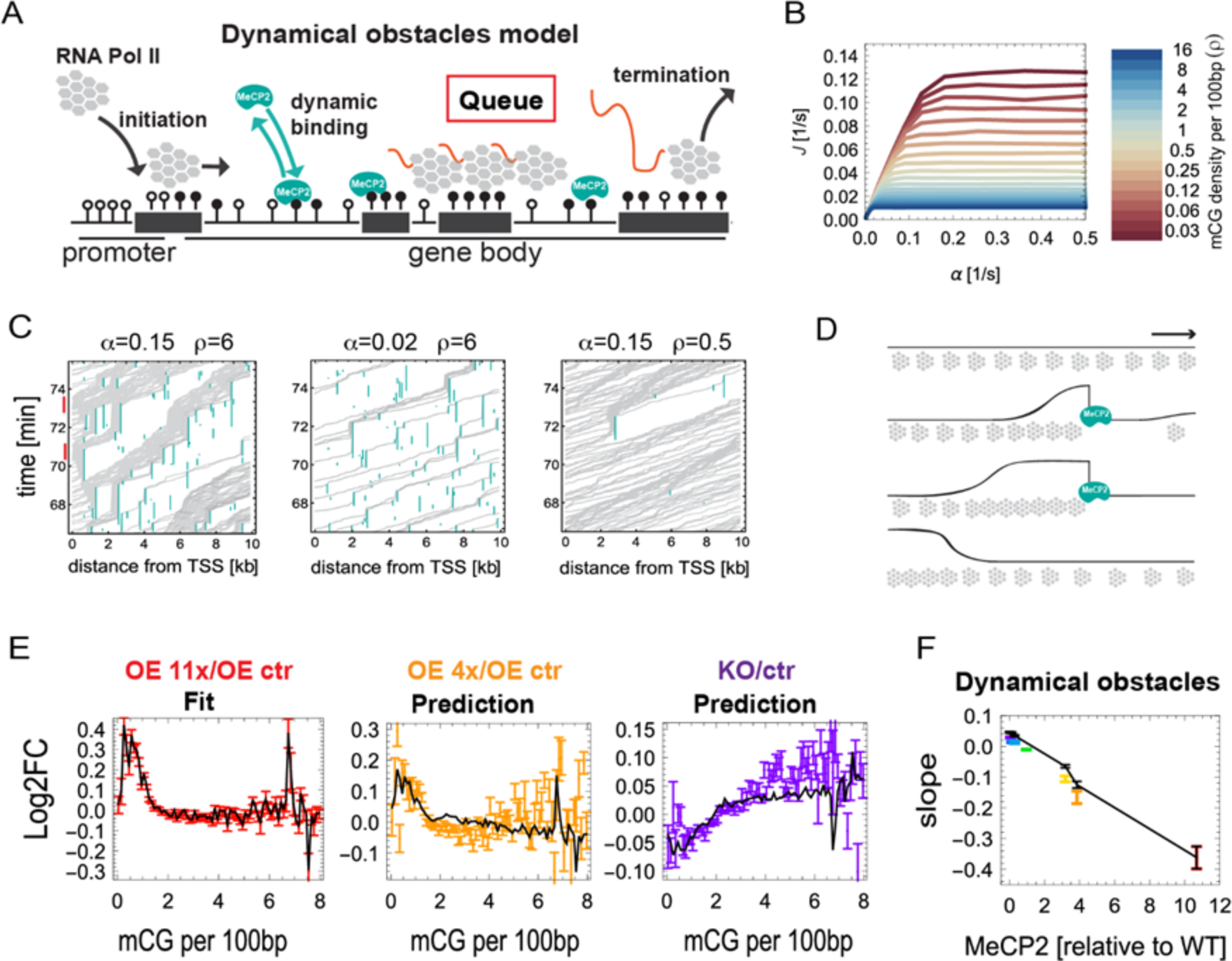
Mathematical modelling shows that MeCP2 slows down transcriptional elongation. (A) Schematic representation of the Dynamical obstacles model. (B) Transcription rate *J* predicted by the model, plotted as a function of the initiation rate *α*, for different gene body mCG densities. (C) Space-time plots (kymographs) representing Pol II moving along the gene. Queues of Pol II induced by MeCP2 can reach TSS (red) and block initiation if both the initiation rate (α) and the density of mCG sites (ρ) are sufficiently high (left panel). (D) Schematic representation of Pol II (grey) density shock waves forming behind MeCP2 (blue). Black line is the local density of Pol II. (E) Log2FC (gene expression) versus mCG density in gene bodies obtained in computer simulations of the Dynamical obstacles model (black solid lines) fitted to the OE 11x/OE ctr RNA-seq dataset (red) agrees well with experimental data for OE 4x/OE ctr (orange) and KO/ctr (purple) datasets. Error bars represent +/− SEM. (F) The maximum slope of Log2FC (gene expression) versus mCG density in gene bodies, predicted by the Dynamical obstacles model (black line). Points are experimental slopes from Fig. 2C.

### MeCP2 binding to both DNA and NCoR are essential to slow down RNA Pol II

To settle the question of whether MeCP2 impedes Pol II progression directly by steric interference or indirectly by altering chromatin structure (e.g., by histone deacetylation (26)), we overexpressed mutated forms of MeCP2 in the presence of WT MeCP2. The mutants were either unable to bind methylated DNA (R111G) (27) or unable to recruit the histone deacetylase complex NCoR (R306C) (14, 28) (Figs. 5A and S8A). As expected, 7-fold overexpression of MeCP2-R111G caused no mCG-density dependent transcriptional changes (Figs. 5B,C and S8B,C). The R306C mutant, on the other hand, was predicted to repress transcription if inhibition is directly due to MeCP2 binding to DNA, but not if inhibition is mediated via the corepressor. In fact, 11-fold overexpression of MeCP2-R306C relative to WT MeCP2 caused only a small perturbation of gene expression, indicating a significant loss of DNA methylation-dependent repression (Figs. 5B,D and S8B,C). The weak slope may represent minor direct interference of DNA-bound MeCP2-R306C with transcription. As neither mutant falls on the line defining the linear relationship between gene repression and MeCP2 concentration (Fig. 5E), our findings favour a predominantly indirect mechanism of repression, whereby corepressor recruitment alters the chromatin state to impede transcription.

**Fig. 5.**
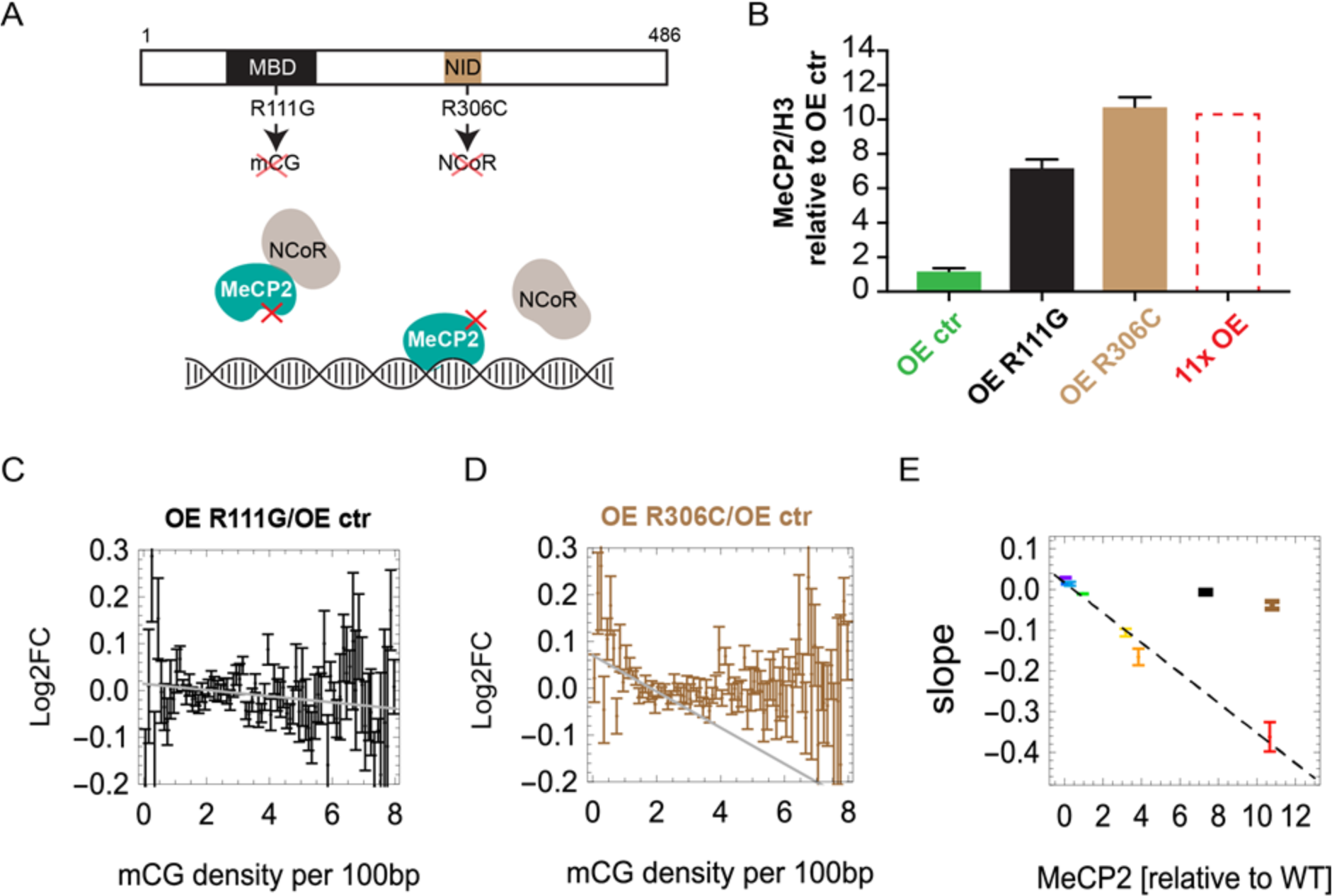
MeCP2 slows down transcription via a mechanism involving NCoR. (A) Location of two binding domains in MeCP2 that are relevant for the proposed mechanism: methyl-CpG binding domain (MBD) and NCoR-interaction domain (NID). The mutation R111G causes MeCP2 to lose the ability to bind specifically to mCG. The mutation R306C prevents MeCP2 from binding the NCoR complex. (B) Level of MeCP2 (Western blot) in two overexpressed mutant cell lines (R111G and R306C) and the overexpression control cell line (OE ctr). OE 11x is shown for comparison. Values are averaged over three biological replicates and normalised by the level of histone H3. (C) Log2FC (expression) of OE R111G/OE ctr shows almost no dependence on mCG density in gene bodies (black). Grey line shows the maximum slope. (D) Log2FC (expression) of OE R306C/OE ctr shows a small negative correlation with gene body mCG density (brown). Grey line shows the maximum slope. (E) Maximum slopes for all cell lines including OE R111G (black) and OE R306C (brown) from (C and D) versus MeCP2 level (Western blot). In all plots error bars represent +/−SEM.

In summary, a close alliance between mathematical modelling and molecular biology has allowed us to discriminate molecular mechanisms underlying the relatively subtle global effects of MeCP2 on global gene expression. The proposed mechanism relies on MeCP2-NCoR interaction that slows down the progression of Pol II during transcription elongation. A candidate mediator of this effect is histone modification, in particular histone deacetylation. The NCoR1 and NCoR2 complexes both recruit histone deacetylases whose action would increase the affinity between DNA and nucleosomes and thus inhibit transcription (26). In fact, cell transfection assays using MeCP2 and methylated reporters have demonstrated that repression depends upon histone deacetylase activity (11, 12).

To explain the dramatic reversibility of Rett syndrome in animal models (29) we propose that, in the absence of MeCP2, DNA methylation patterns are unaffected, allowing the re-expressed wildtype protein to bind within gene bodies and commence normal modulation of transcriptional elongation. The Congestion Model may apply to proteins other than MeCP2. For example, other chromatin-binding factors that bind short (and thus abundant) motifs, including other methyl-binding proteins, may modulate gene expression by a similar mechanism.

## Materials and methods

### Cell lines

The procedure for culture and differentiation of the LUHMES cell line was previously described (18). To create two independent *MECP2* knock-out lines, we used CRISPR-mediated gene disruption (30). To generate MeCP2 knock-downs, several shRNAs against MeCP2 were designed using Sigma-Aldrich Mission shRNA online software. Two shRNAs were chosen and cloned into pLKO.1 vector including scrambled shRNA as a control and lentiviruses were created (Table S2). To increase the level of MeCP2 we created lentiviruses expressing MeCP2 from two alternative promoters in the pLKO.1 vector: Synapsin and cytomegalovirus (CMV). The human Synapsin promoter drove expression of the MeCP2 E2 isoform, while the CMV promoter drove the MeCP2 E1 isoform. As a control, we expressed GFP under the control of the CMV promoter. Optical and immunofluorescence microscopy was used to confirm differentiation into neurons. Western blot was used to quantify the level of MeCP2. Global methylation levels were quantified using HPLC on the 5 μm Apex ODS C18 column, with isocratic 50 mM ammonium phospate (monobasic) mobile phase. Calculation of standard deviation, standard error of mean and *t* tests for qPCR, Western blots, methylation and total RNA quantification using HPLC were performed using GraphPad Prism version 7. See SI Methods Section 1 for details.

### Repression assay

CpG-free vector containing Firefly Luciferase with CpGs was methylated by M.SssI methyltransferase in presence or absence of SAM. Mouse embryonic fibroblasts were transfected using Lipofectamine 2000 with three plasmids containing: Firefly Luciferase, Renilla Luciferase and MeCP2. Luciferase activity measurements were performed using Dual Luciferase assay kit (Promega) according to manufacturer protocol. See SI Methods Section 1.6 for details.

### Library preparation for Illumina sequencing

All libraries were sequenced as 75- or 100-nucleotide long paired-end reads on HiSeq 2000 and HiSeq 2500 Illumina platforms. Methylome of wildtype LUHMES-derived neurons at day 9 was obtained by TAB-seq according to the published protocol (31). RNA-seq library was performed according to manufacturer protocol for ScriptSeq Complete Gold kit (Human/Mouse/Rat). Total RNA was isolated from all generated cell lines (Table S1) at day 9 of differentiation using either the RNeasy Mini kit or the AllPrep DNA/RNA Mini kit (Qiagen). ATAC-seq in four cell lines (KO, WT, OE 4x and OE 11x, see Table S1) was performed as in (32). See SI Methods Section 2 for details.

MeCP2 ChIP-seq was performed using LUHMES-derived neurons at day 9 of differentiation with four levels of MeCP2: KO, WT, OE 4x and OE 11x (see Table S1). Cells were crosslinked with 1% of Formaldehyde (Sigma) in the medium for 10 min at RT and quenched with 2.5 M Glycine (Sigma) for 2 min at RT. Crosslinked and sonicated chromatin was mixed with 60 ng of sonicated *Drosophila* chromatin (Active Motif) as a spike-in, and the mix was incubated overnight at 4 ºC with antibodies against MeCP2 (D4F3, Cell Signalling) plus spike-in antibody (Active Motif). For ChIP-seq library preparation, IPs for each condition were pooled together to achieve 5 ng total DNA as a starting material. Libraries were prepared using the NEBNext Ultra II DNA library Prep kit (NEB) for both IPs and corresponding inputs. See SI Methods Section 2.4 for details.

### Data processing of raw reads from Illumina sequencing

All reads were quality-controlled, trimmed to remove adapters (Trimmomatic) (33), and duplicated reads, and mapped to the human hg19 reference genome. Bismark (34) was used to extract cytosine methylation from TAB-seq. Total methylation (total mCG) in a gene was calculated as the total number of methylated CG dinucleotides. mCG density was calculated by dividing total methylation by the length of the corresponding region (gene body or promoter). mCG mean was defined as the total mCG divided by the number of all CG dinucleotides and multiplied by 100. See SI Methods Section 3 for details.

### ChIP-seq enrichment profiles

We first obtain accumulated counts (the number of reads) 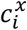 that overlap with *i*-th basepair to the right (*i* > 0) or left (*i* < 0) from feature *x* (*x* =mCG, mCA, …). We then calculate enrichment profiles as

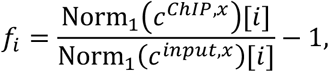

where 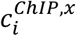 and 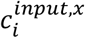 are accumulated counts from ChIP and input (genomic) DNA sequencing, respectively, and Norm_1_(*c*)[*i*] normalizes the counts profiles such that their flanks have values close to one:

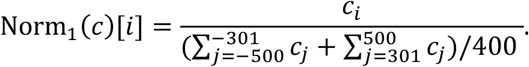

We consider a particular C to be methylated if it is methylated in 100% of the reads, and the coverage is at least 5. We consider a C to be unmethylated if it does not show up in any of the ChIP-seq reads as methylated. See SI Methods Section 4 for details.

### Computer model of ChIP-seq

We assume that MeCP2 occupies methylated cytosines with probability *p* times the probability of binding to a particular motif (SI Section 5). Binding probabilities for different motifs are based on known binding affinities (35) and relative binding strengths (15). To create simulated ChIP fragments, we assume that if a DNA fragment contains at least one MeCP2 bound to it, it will be present in the simulated ChIP-seq. Fragments that do not contain any MeCP2 may still be present in the ChIP-seq data with probability *p*_*bg*_ which accounts for “background” reads in ChIP-seq even in the absence of MeCP2. This is similar to previous models of ChIP-seq (36); even best ChIP-seq libraries can have a significant level of noise (*p*_*bg*_ close to 1) (37). We also add CG- and length bias (SI Section 5.1), and process simulated reads in the same way as the experimental ChIP data. For each ChIP-seq data set we fitted the simulated profile (parametrized by *p*, *p*_*bg*_) to the experimental profile. Any *p* ≤ 0.1 gives a good fit (Fig. S5D), indicating that *p* ≈ 0.1 is the upper bound on mCG occupancy in 11x OE. We used best-fit parameters to predict profiles on features other than mCG (Fig. S5E,F). See SI Methods Section 5.3 for details.

### ATAC-seq footprints

ATAC-seq was analysed in a similar way to ChIP-seq, except that we used fragments’ endpoints (Tn5 insertion sites) to generate accumulated counts *n*_*i*_. We calculated the insertion profiles as

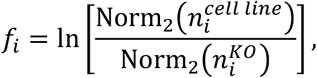

where 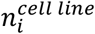 and 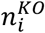 are the insertion counts profiles for a given cell line and KO1,respectively, and Norm_2_ normalizes the counts profiles such that their flanks have values close to one:

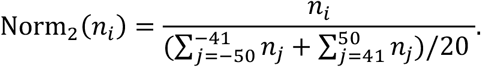

### Computer model of ATAC-seq

We use the same binding model as in the ChIP-seq simulations. We assume that MeCP2 occupies 11bp (20) and that the protein is centred on an mC. We simulate the action of the Tn5 transposase by splitting the sequence into fragments in areas free of MeCP2, and we include Tn5 sequence bias, and CG- and length bias (SI Section 6). The model has three parameters: the density *p* of MeCP2 on mCxx, the average density of insertion (cut) sites *t*, and the GC bias *b*. We process simulated DNA fragments in the same way as described above for the experimental data. We examined the role of the parameters on the shape and depth of the simulated footprint of MeCP2 (SI Section 6.2) and concluded that the footprint is not affected as long as the test and control samples have been processed in a similar way. To extract MeCP2 occupancy *p* from ATAC-seq data, we fitted the model (free parameters *p*, *t*, and a fixed *b* = 6.0) to experimental footprints for all four cell lines. The relationship is linear (Fig. 2E), with the best-fit *p* = 0.0058 × *M*_*cell line*_/*M*_*WT*_. See SI Methods Section 6 for details.

### Chromatin accessibility from ATAC-seq

For each gene, we calculated its mean insertion count *n̅* and selected regions (“insertion peaks”) in which *n*_*i*_ > 4*n̅*. Accessibility was defined as the sum of all insertions in the peaks divided by the “background” *n̅*:

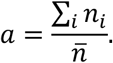

### RNA-seq data analysis

We used a subset of protein-coding genes with sufficient methylation coverage (BS-seq; ≥80% C detected as methylated, coverage ≥20), and gene bodies 1kb or longer. This resulted in 15382 genes out of the initial 17764 protein-coding genes (86%).

To obtain the slope (Fig. 1C) we fitted straight lines Log2FC = *aρ*_mCG_ + *b* to Log2FC(expression) to different ranges of *ρ*_mCG_ ∈ [*ρ*_1_, *ρ*_1_ + Δ*ρ*], for different pairs *ρ*_1_ ∈ [0.5,1] and Δ*ρ* ∈ [1,4] (units: 1/100bp). The maximum slope equals *a* of the fit with the largest absolute slope to error ratio (largest |*a*|/*σ*_*a*_ where *σ*_*a*_ is the standard error of *a*).

In all plots of Log2FC of differential gene expression we shifted the Log2FC values so that the average Log2FC in the range of mCG density *ρ*_mCG_ ∈ [1,6] bp/100bp was zero for all samples. This was motivated by a difficulty in determining the absolute levels of expression since we did not quantify total mRNA.

### Mathematical models of gene expression

The condensation model assumes that the fraction *f*_*i*_ of cells in which gene *i* is actively transcribed depends on promoter openness *a*_*i*_ (measured by ATAC-seq) which in turn depends on the level *M* of MeCP2 and gene methylation *ρ*_*i*_: *f*_*i*_ = *f*_*i*_ (*M*, *ρ*_*i*_) ∝ *a*_*i*_ = *a*_*i*_(*M*, *ρ*_*i*_). The model predicts (SI Section 8) that Log2FC_X/Y_ of the ratio of gene expression of cell line X versus cell line Y should yield the same curve (plus a constant) as the logarithm of the ratio of accessibilities of X versus Y when plotted as a function of *ρ*_mCG_. Data does not support this model (Fig. 3C). The detachment model poses that the probability that RNA Pol II successfully terminates is *P* = (1 − *λ*)^*n*^ ≅ *e*^−*λn*^, where *n* is the number of “abort sites” on the gene, proportional to the number of MeCP2 molecules on the gene, and *λ* is the abortion probability. We show (SI Section 9) that

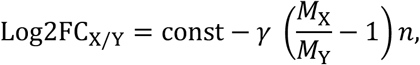

where *γ* ∝ *λ* is an unknown parameter identical for all cell lines, and *M*_X_, *M*_Y_ are MeCP2 levels in cell lines X and Y. The model is rejected (Fig. 3E).

We consider two mechanisms by which MeCP2 could affect elongation. To implement the slow sites model we use the totally asymmetric simple exclusion process (TASEP) with open boundaries (24). A gene is represented as a chain of *L* sites. Each site (equivalent to 60bp of the DNA) is either occupied by a particle (RNA Pol II) or is empty. Particles enter the chain at site *i* = 1 with rate *α* (transcription initiation rate), move along the chain and exit at site *i* = *L* with rate *β* = 1 sec^−1^. Sites can be “fast” or “slow”. Slow sites represent mCGs affected by the interaction with MeCP2, whereas fast sites are all other sites (methylated or not). Particles jump with rate *v* = 1 sec^−1^(equivalent to Pol II speed ≈60bp/s) on fast sites and *v*_*s*_ = 0.05 sec^−1^ on slow sites. Slow sites are randomly and uniformly distributed with density *ρ*_*s*_ = *pρ*_mCG_ where *p* is the probability that an mCG is occupied by MeCP2. See SI Section 10.2 for more details. To relate this model to the mRNA-seq differential expression data we calculate Log2FC as

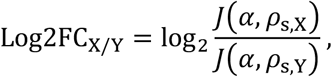

where *ρ*_s,X_ = *ρ*_mCG_*p*_X_, *ρ*_s,Y_ = *ρ*_mCG_*p*_Y_ in which *p*_X_, *p*_Y_ are MeCP2 occupation probabilities for cell lines X,Y. In the above expression we know all quantities except the initiation rate *α* which we fit to the OE 11x data (SI Section 10).

The dynamical obstacles model is very similar with two exceptions: (i) Pol II always moves with the same speed *v* (no slow sites) as long as it is not blocked by other polymerases and obstacles, (ii) obstacles bind and unbind dynamically from the methylated sites. We assume that unbinding occurs with rate *k*_*u*_ per obstacle, whereas binding occurs with rate *k*_*u*_*p* per unoccupied mCG. Obstacles do not bind if an mCG is already occupied by an obstacle or a polymerase. We assume that obstacles are not restricted to accessible mCGs and that their density on actively transcribed genes may be higher than *p* obtained from ATAC-seq but still proportional to MeCP2 level. We found that *p* = *M*/*M*_*0E11x*_ reproduces Log2FC data for all cell lines. See SI Methods Section 10.3 for details.

## Supporting information

Table S4: q-PCR primers used in this work.

Table S3: The list of antibodies used in this work.

Table S2: Sequences of shRNA and guide RNA used to make knock-downs and knock-outs.

Table S1: Cell lines, their MeCP2 levels, and number of replicates for each experiment (RNA-seq, ATAC-seq, ChIP-seq)

SI Video 2: Congestion model with dynamic obstacles: visualisation of Pol II traffic in the presence of MeCP2-induced obstacles (blue rectangles).

SI Video 1: Congestion model with dynamic obstacles: visualisation of Pol II traffic in the absence of obstacles.

SI Video 3: Congestion model with slow sites: visualisation of Pol II traffic in the presence of MeCP2-induced slow sites.

## Acknowledgments

We thank Tanja Waldmann for introducing us to the LUHMES cell line, Beatrice Alexander-Howden for technical support, David Kelly for microscope assistance, Martin Waterfall for help with FACS and Jim Selfridge for help with preparing samples for HPLC. We thank Sabine Lagger and John Connelly for critical assessment of the manuscript. The work has made use of resources provided by the Edinburgh Compute and Data Facility (ECDF; www.ecdf.ed.ac.uk) and was supported by a Wellcome Trust Programme Grant and Investigator Award to AB. JCW was supported by a grant from the Rett Syndrome Research Trust. RS was supported by a Wellcome Trust 4 year PhD studentship. BW was supported by a Personal Research Fellowship from the Royal Society of Edinburgh.

## Author contributions

J.C.W and R.S. designed experiments, made and characterised cell lines and interpreted data; J.C.W performed the NGS experiments; S.W and K.C. performed NGS data preparation; B.R. performed HPLC analysis; M.Y. performed initial reaction for TAB-seq; P.G. and B.W. developed the “Congestion model”. B.W. conceived the mathematical modelling, analysed NGS data and interpreted data. A.B. conceived the study and, together with J.C.W., R.S. and B.W., wrote and edited manuscript.

---

**Fig. S1.**
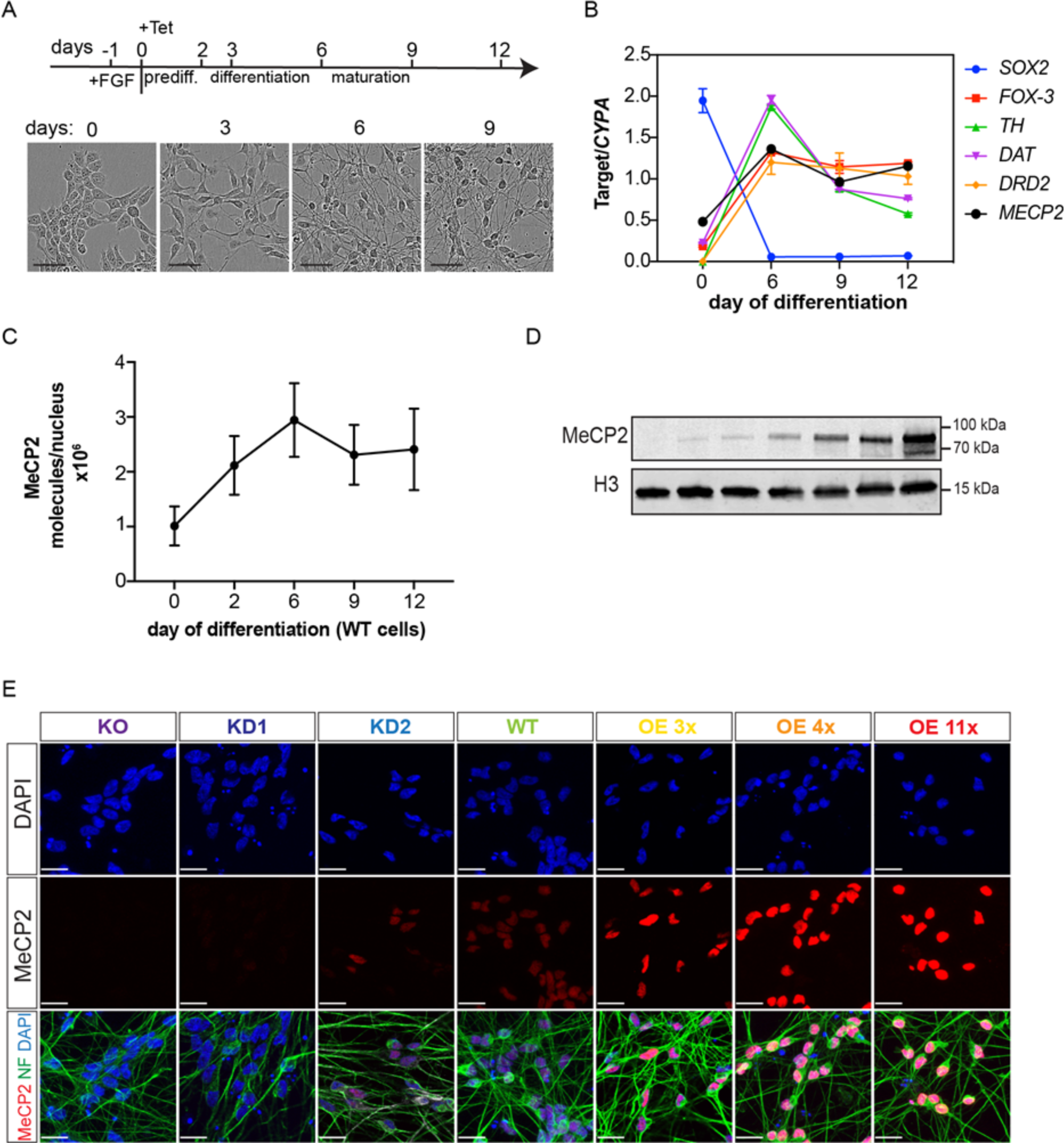
Characterisation of LUHMES-derived cell lines KO and WT. (A) Experimental protocol for LUHMES differentiation. Phase contrast images show cells in different stages of differentiation. Scale bar is 50 μm. (B) Expression of neuronal differentiation markers normalized to the housekeeping gene *CYPA* during WT LUHMES differentiation. Error bars represent +/−SEM. (C) Changes in the number of MeCP2 molecules per nucleus during differentiation of WT LUHMES cells, calculated from Western blotting. Error bars represent +/− SEM. (D) Expression of MeCP2 in cells at day 9 of differentiation (Western blot). (E) Immunofluorescence imaging of MeCP2 and neurofilament (NF) expression in day-9 neurons. Scale bar is 20 μm.

**Figure S2.**
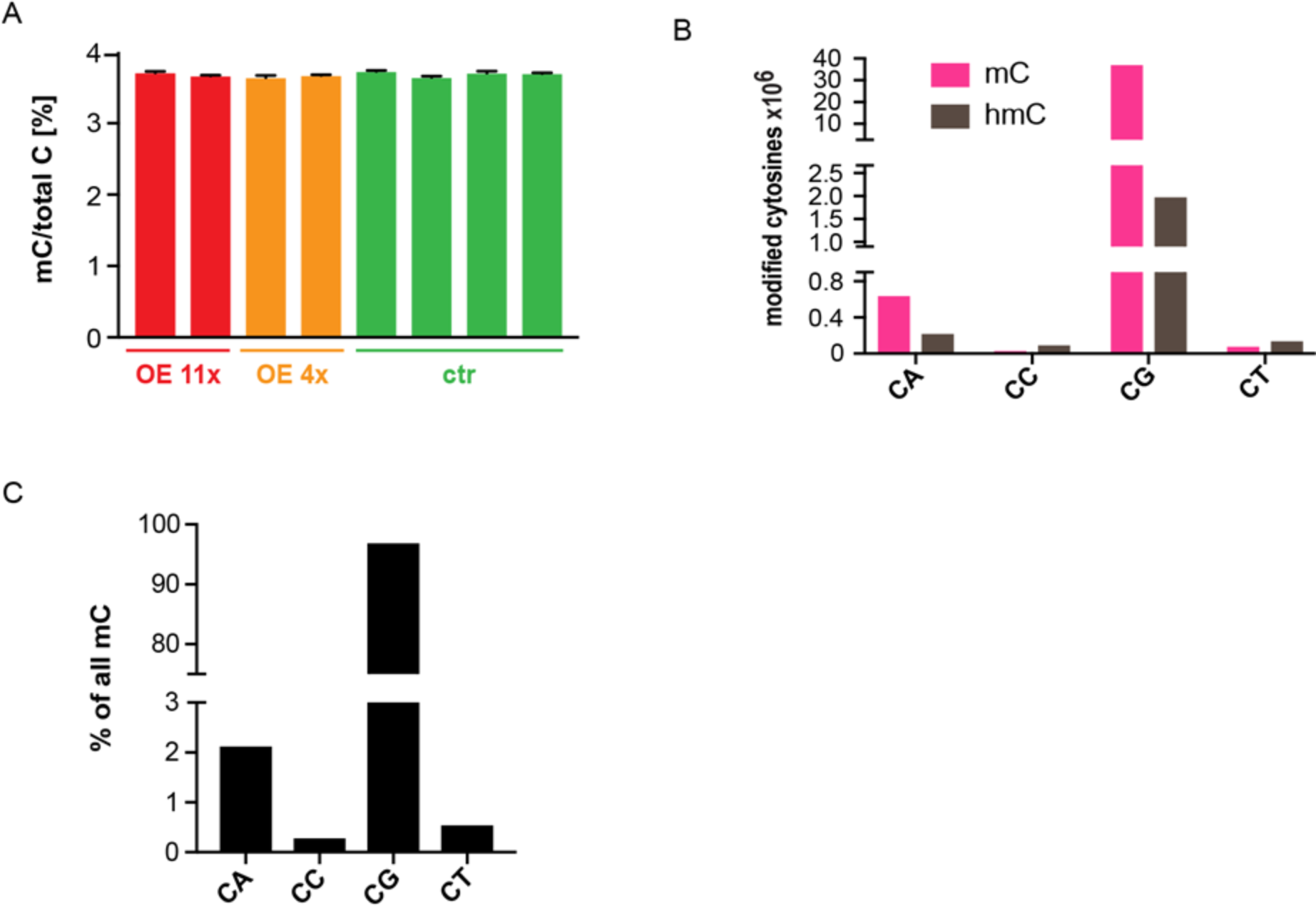
DNA methylation in LUHMES WT cells. (A) Fraction of methylated Cs in the genome quantified by HPLC for LUHMES-derived neurons at day 9 of differentiation. OE 11x (red), OE 4x (orange) and controls (green). Error bars are +/−SEM. (B) Number of methylated (mC) and hydroxymethylated (hmC) dinucleotides (per haploid genome) obtained from TAB-seq in WT LUHMES-derived neurons (day 9). (C) Percentage fraction of methylated Cs in the context of different dinucleotides calculated from TAB-seq data.

**Figure S3.**
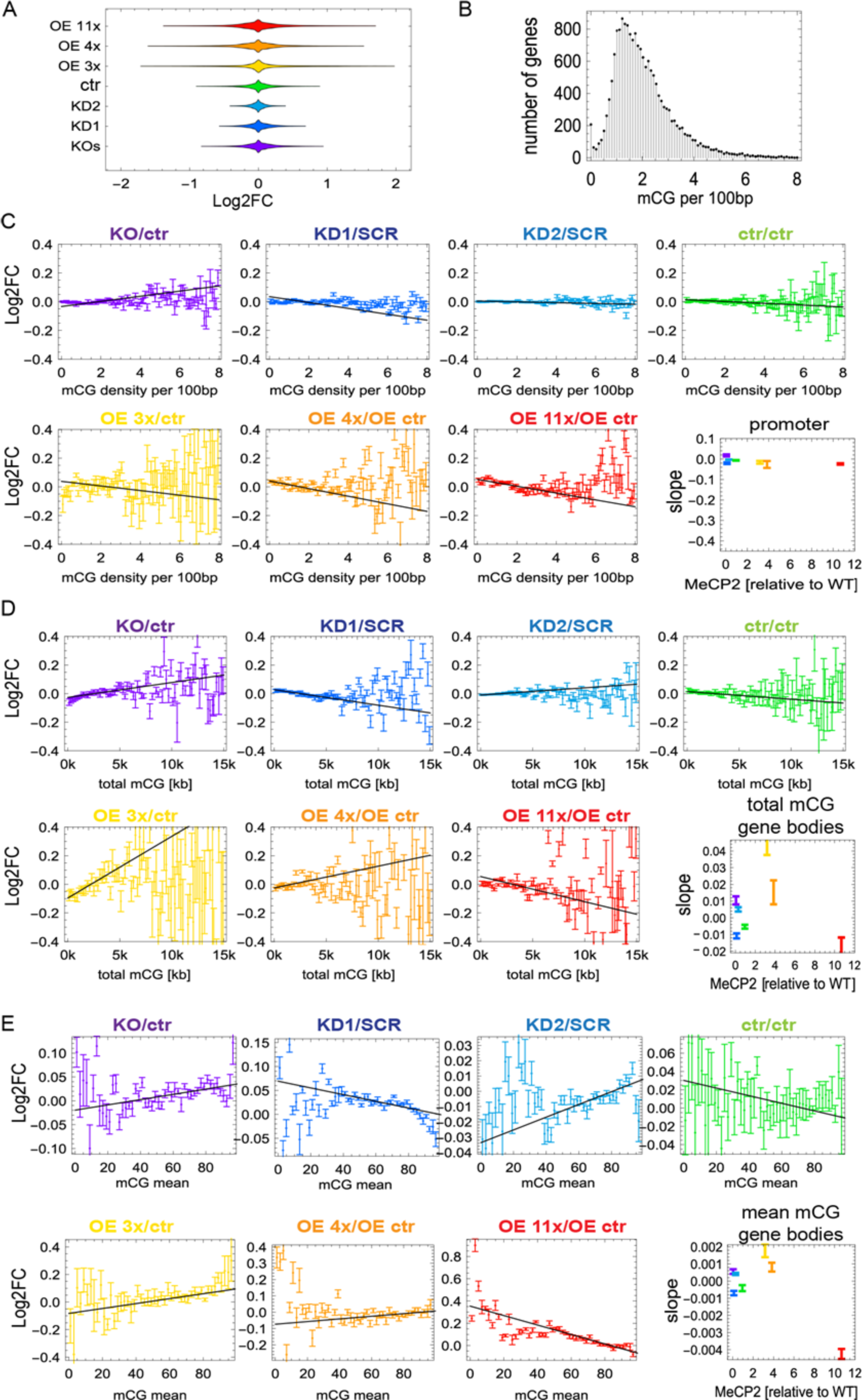
Gene expression changes do not correlate with other methylation-related quantities such as total mCG and mCG mean. (A) Violin plots show that changes in gene expression in cell lines expressing different levels of MeCP2 are very small. (B) Number of genes versus mCG density in gene bodies. Genes have been binned as in Fig. 1C (bin width 0.1bp). (C) Log2FC of gene expression relative to appropriate controls (ctr – unmodified controls; SCR – scrambled control, OE ctr – overexpression control) for all seven levels of MeCP2, plotted against mCG density at promoters. Genes were binned according to their promoter mCG density (bin size = 0.1bp), with each point representing a mean Log2FC of all genes falling in that particular bin. Black line shows the maximum slope. The slope of Log2FC for promoter mCG shows minimal dependence on the level of MeCP2. (D) As in C but for total mCG. The slope of Log2FC does not show a clear dependence on the level of MeCP2. (E) As in C but for mCG mean. In all panels, error bars represent +/− SEM.

**Fig. S4.**
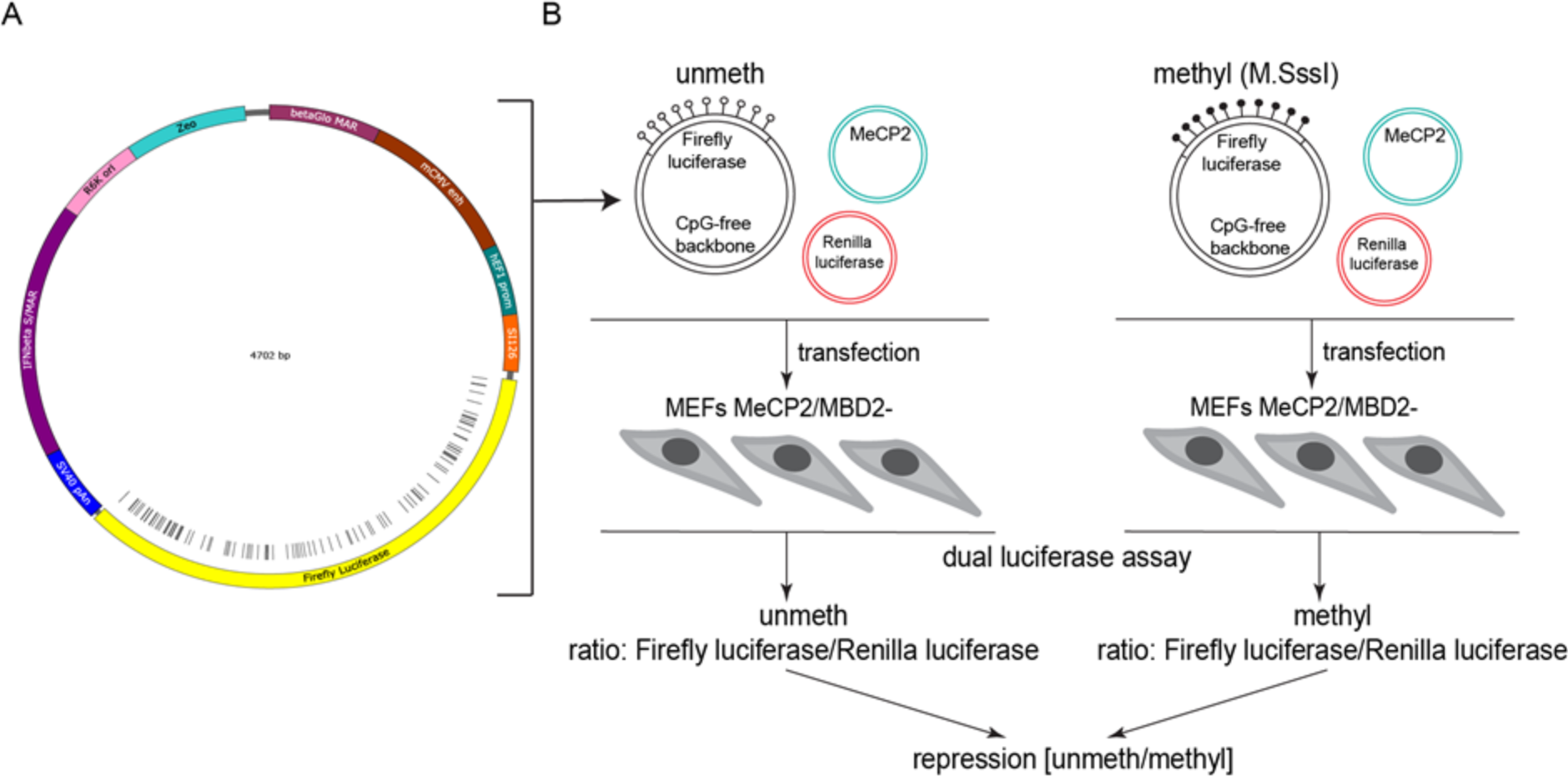
Repression assay designed to measure MeCP2-mediated repression. (A) A map of the CpG-free vector used to express luciferase containing approximately 100 CpGs restricted to the body of the luciferase gene. (B) Study design shows how the expression of un-methylated and methylated reporter are compared, each normalised to a co-transfected construct expressing Renilla luciferase. Transfected mouse fibroblasts cells were null for both *Mecp2* and *Mbd2* genes.

**Figure S5.**
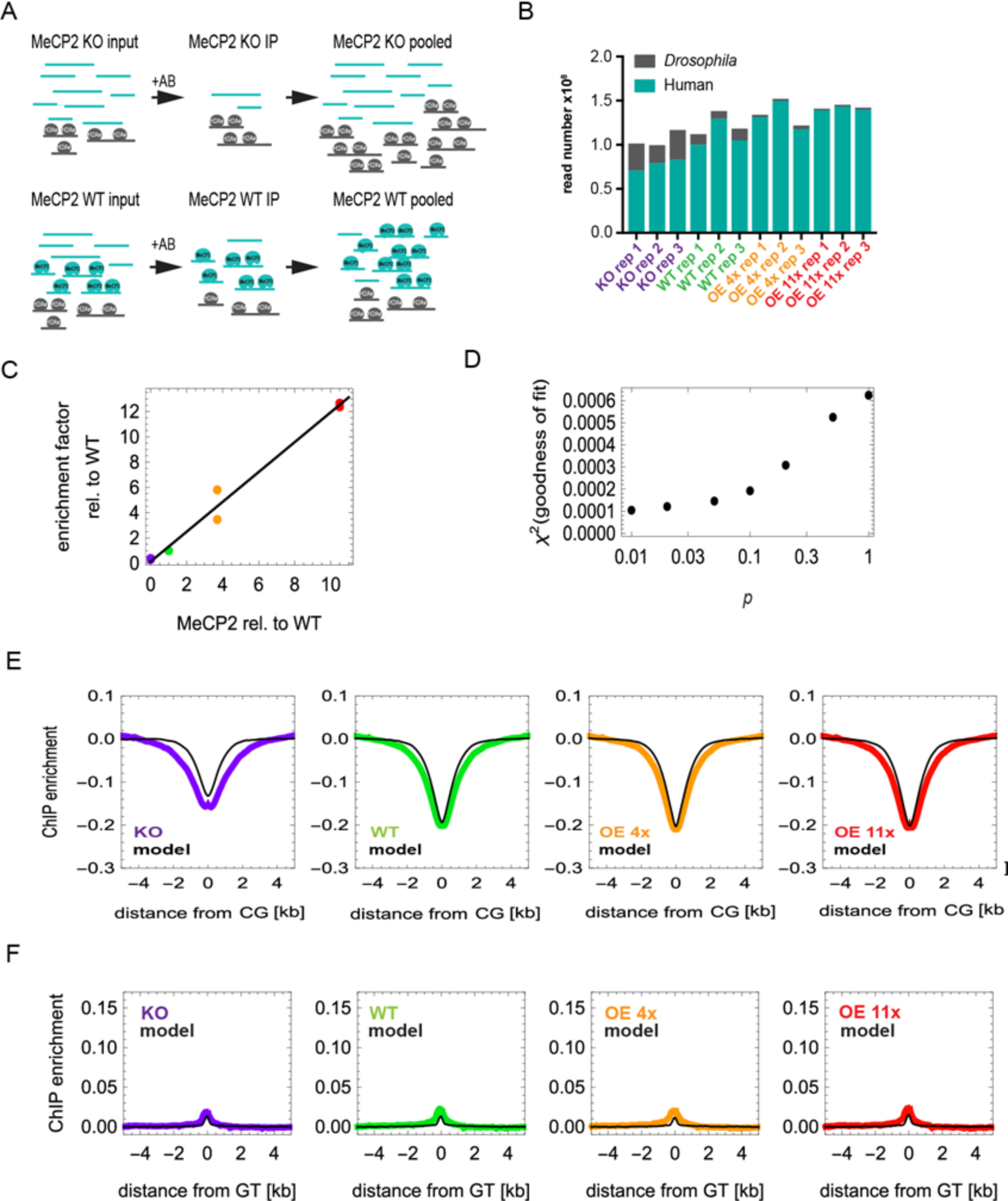
Details of the experimental and simulated ChIP-seq. (A) Schematic representation of our quantitative MeCP2 ChIP-seq protocol for human neuronal chromatin (turquoise), using Drosophila chromatin (grey) as a spike-in for normalisation, precipitated using antibodies against H2Av. (B) Total number of *Drosophila* reads compared to LUHMES reads obtained from ChIP-seq, for four MeCP2 levels: KO (purple), WT (green), OE 4x (orange) and OE 11x (red); each with three biological replicates. (C) Enrichment factor (human chromatin relative to spiked-in *Drosophila* chromatin) increases linearly with the level of MeCP2. Two biological replicates are shown as individual data points. (D) ChIP-seq model goodness-of-fit versus *p* (probability that an mCG is occupied by MeCP2). The model becomes relatively insensitive to the exact value of *p* for *p* < 0.1. (E) ChIP-seq enrichment profiles centred at unmethylated CG dinucleotides. Black lines show profiles predicted by the model. (F) ChIP enrichment profiles on unmethylated GT. The model (black lines) correctly predicts the lack of enrichment.

**Fig. S6.**
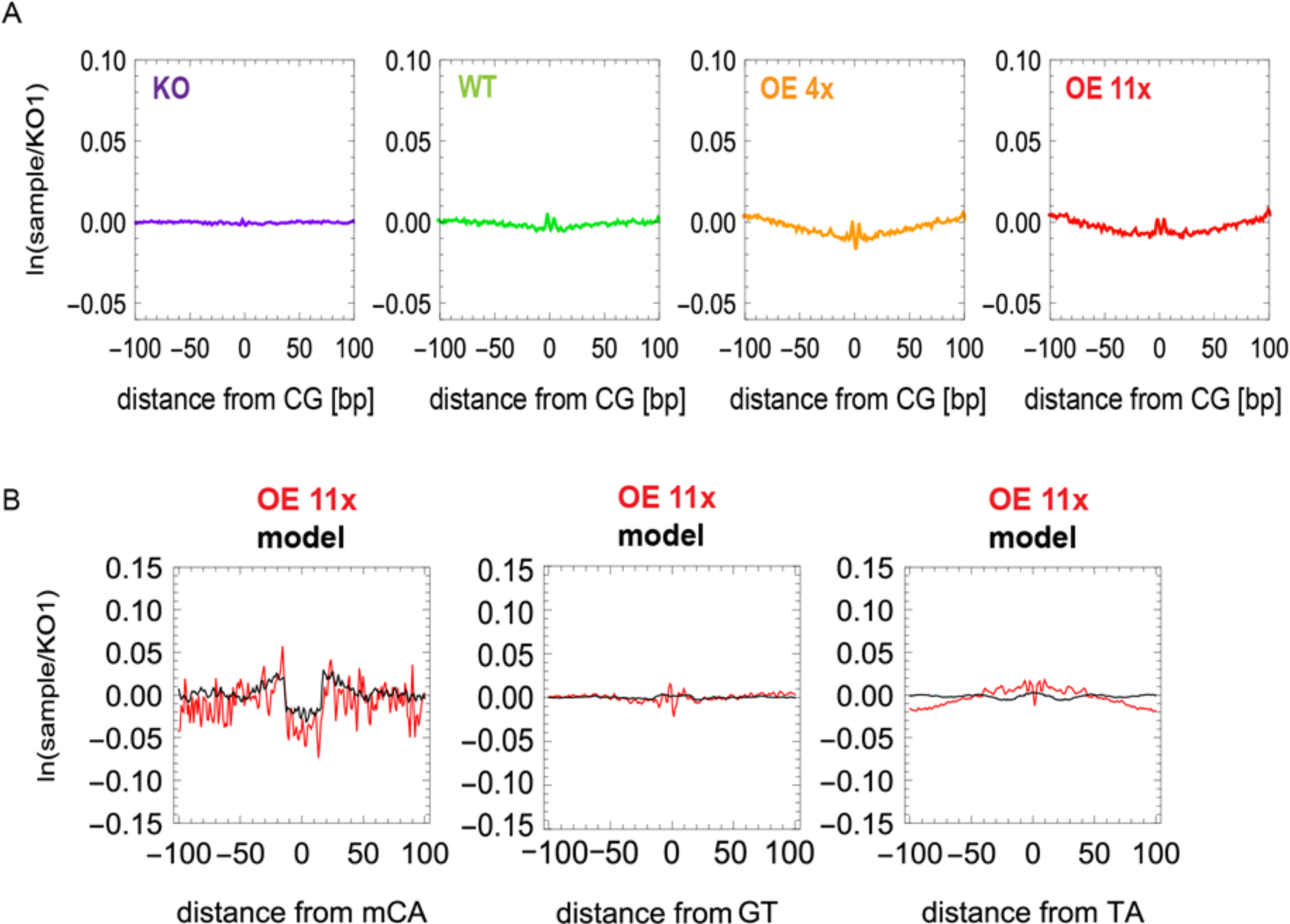
MeCP2 footprint is found on mCA but not on unmethylated dinucleotides. (A) ATAC-seq depletion profiles in the +/−100 bp regions surrounding unmethylated CG dinucleotides. Profiles are averages over 2-4 biological replicates. (B) MeCP2 footprint on features other than mCG: mCA, GT, TA for OE 11x (red) normalized by KO, compared to predicted footprint profiles from simulations (black). As in ChIP-seq, the footprint is present on mCA and absent on GT and TA.

**Figure S7.**
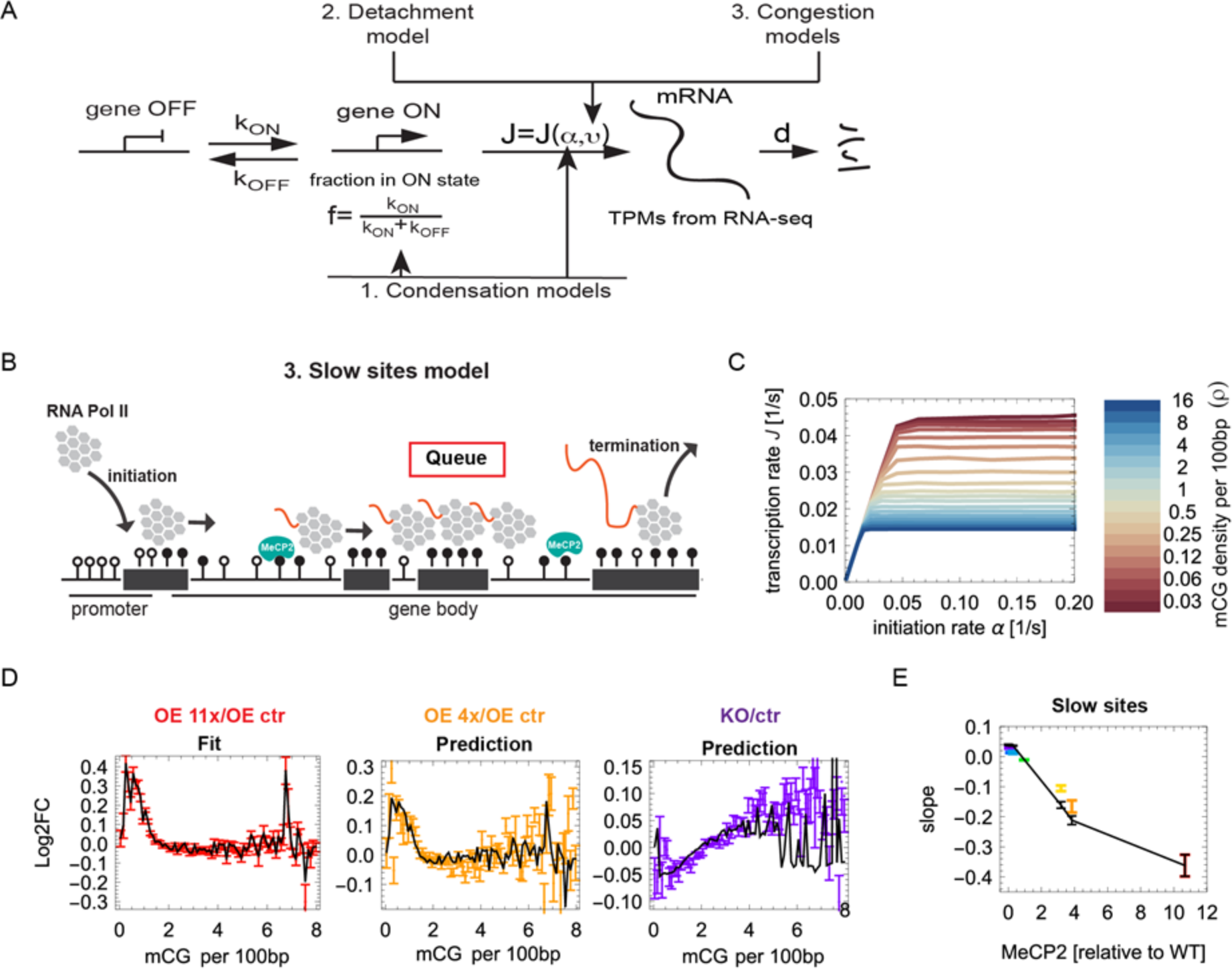
Slow Sites model. (A) A general model of gene transcription. Each gene can be in two states: ON (TSS accessible to Pol II) or OFF (TSS not accessible). In the ON state, mRNA is produced with rate *J* which depends on the transcription initiation rate *α* and the elongation rate *v*. RNA-seq does not directly measure *J* but it gives the amount of mRNA accumulated in the cell (TPMs, transcripts per million bp) which also depends on degradation rate *d*. Three proposed models of MeCP2-dependent transcriptional regulation relate to different stages of transcription. (B) The Slow Sites model in which MeCP2-induced chromatin modifications slow down elongating RNA Pol II. (C) Transcription rate *J* as a function of the initiation rate α. *J* saturates for high *α*, similarly as in the Dynamical Obstacles model (Fig. 4B). Log2FC obtained from the Slow Sites model (black line) agree well with experimental data (red, orange and purple). Left panel: model fitted to OE 11x to obtain *α(ρ)*. Middle and right panel: model predictions versus experimental Log2FC for OE 4x (orange) and KO (purple). The maximum slope of Log2FC (gene expression) versus mCG density in gene bodies as predicted by the Slow Sites model (black line) reproduces the slopes from RNA-seq data for all seven levels of MeCP2.

**Fig. S8.**
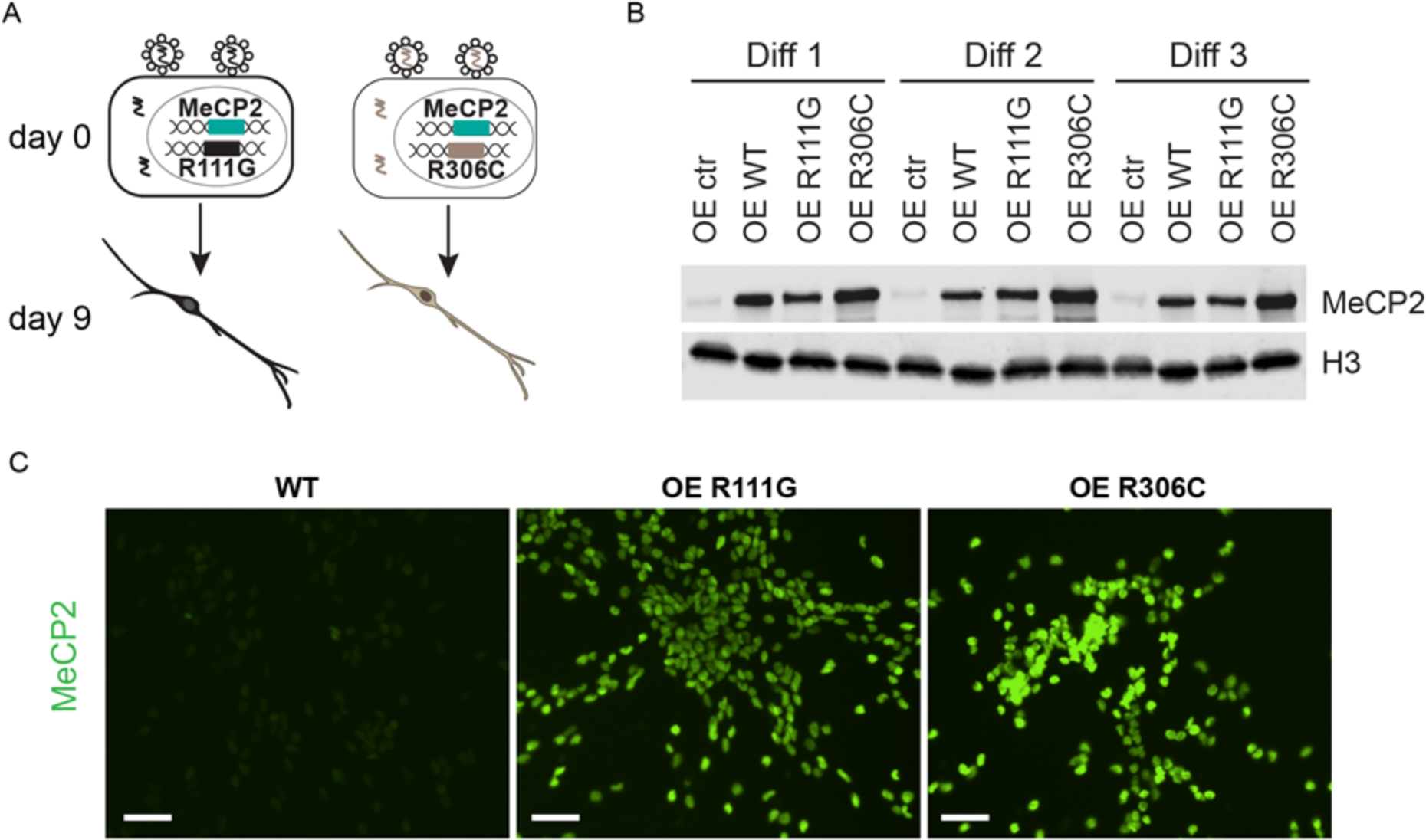
Generation of MeCP2 overexpression mutants. (A) LUHMES cell lines were modified to overexpress MeCP2 with different mutations R111G (black) and R306C (brown). (B) Overexpression of R111G and R306C compared with control cells (OE ctr) and OE WT (11x) confirmed by Western blots using antibodies against MeCP2 and H3 as loading control in three independent differentiations. (C) Immunofluorescence images with an antibody against MeCP2 show uniform overexpression of MeCP2 mutants R111G and R306C in the LUHMES-derived neurons at 9 days of differentiation. Scale bar is 50 μm.

Table S2: Sequences of shRNA used to make knock-downs.

Table S3: The list of antibodies used in this work.

Table S4: q-PCR primers used in this work.

