## Supplementary information for "Quantitative modelling predicts the impact of DNA methylation on RNA polymerase II traffic"

#### 1. Cell lines and experimental methods.

##### 1.1. LUHMES culture and differentiation.

Proliferating LUHMES cells were cultured on Nunc plasticware coated with 40 µg/ml Poly-L-ornithine (Sigma-Aldrich; P-3655-100mg) and 1µg/ml Fibronectin (Sigma-Aldrich; F-1141-5mg) in medium for proliferating cells: Advanced DMEM (Life Technologies; 12634-010) plus N2 supplement (Life Technologies; 17502-048), 2mM L-Glutamine (Sigma-Aldrich; G7513), and 40 µg/ml bFGF (R&D Systems; 4114-TC-1 mg). For differentiation, LUHMES cells were seeded on 40 µg/ml Poly-L-Ornithine and 1 µg/ml Fibronectin coated plastic-ware at a density of 3-4 x $10^4$ cells/$cm^2$ in medium for proliferating cells. Next day after seeding, the medium was changed to the differentiation medium containing Advanced DMEM, 2mM L-Glutamine, N2 supplement, 1mM cAMP (Sigma-Aldrich; D0627), 1µg/ml Tetracycline (Sigma-Aldrich; T-7660) and 2 ng/ml GDNF (R&D Systems; 212-GD-50 µg). Two days later, cells were trypsinised and seeded at a density of 1.1x$10^5$ cells/$cm^2$ in the differentiation medium for the final differentiation (Fig. S1A).

##### 1.2. Lentiviral particle preparation and LUHMES infection.

Lentiviruses were produced in HEK293FT cells (Life Technologies; R700-07) by retrograde transfection of vectors: 7.5 µg pLKO.1; 4.6 µg psPax2; 2.8 µg pMD2G using Lipofectamine 2000 (Life Technologies; 11668027). For one transfection, plasmids in appropriate ratios were added to 1.5 ml OptiMEM (Life Technologies; 31985062) and 36 µl of Lipofectamine 2000 was mixed with 1.5 ml of OptiMEM. Both solutions were combined and incubated for 20 min at room temperature. Meanwhile, HEK293FT cells were trypsinised and concentration-adjusted to 1.2x$10^6$/ml. 5 ml of cell suspension was added to a 10 cm dish; the transfection mix was then added and the culture was left overnight. Cells were left for 72 hours, after which the supernatant was collected, and viral particles were precipitated using a PEG Virus Precipitation kit (BioVision) following the manufacturer's protocol. Viruses were aliquoted and either used directly or frozen at -80ºC. LUHMES cells were infected with lentiviral particles overnight. The next day, cells were selected with 0.5 µg/ml Puromycin (Life Technologies; A11138-03), and concentration of antibiotic was reduced the following day to 0.25 µg/ml. The population of resistant cells was either analysed as a pool or FACS-sorted into a 96-well plate in order to isolate single clones.

##### 1.3. Western blot, qPCR, immunofluorescence, live cell imaging.

For protein quantification, cells were either lysed to give a whole cell extract or only nuclei were isolated, depending on the location protein of interest. To obtain whole cell extracts, cells were homogenized in the NE1 buffer (20 mM HEPES pH 7.9, 10 mM KCl, 1 mM $MgCl_2$, 0.5 mM DTT, 0.1% Triton X, 20% glycerol, 2mM PMSF, protease inhibitor) and treated with Benzonase (Sigma) for 15 min at RT to remove DNA. Protein concentration was measured (Bradford, Bio-Rad) and loading buffer (5x, 250 mM Tris·HCl pH 6.8, 10% SDS, 30% [v/v] Glycerol, 10 mM DTT, 0.05% [w/v] Bromophenol Blue) was added. Samples were boiled for 5 min prior to loading 40 µg of protein extract onto a polyacrylamide gel.

To prepare nuclei, cells were lysed in hypotonic buffer (0.32 M Sucrose, 10 mM Tris-HCl pH 8, 3 mM $CaCl_2$, 2 mM MgOAc, 0.1 mM EDTA, 0.5% NP-40). Nuclei were washed with NP-40-free buffer, counted, centrifuged, and resuspended in an appropriate volume of 20% glycerol in PBS. Nuclei were treated with Benzonase, loading buffer was added and samples were boiled for 5 min. Nuclear lysates ($1.6x10^5$ per lane) were separated on the gradient 4-15% SDS-PAGE gels (Mini-Protean, Bio-Rad) in running buffer (25 mM Tris, 190 mM glycine, 0.1% SDS) at 200 V for ~40 min alongside a protein size marker (PageRuler, Thermo Scientific). Proteins were transferred onto a Nitrocellulose membrane using wet transfer method for 1 hour at 200V. The membrane was stained first with Ponceau S solution for quality check and then treated with blocking buffer (PBS, 1% PVP, 1% non-fat dried milk, 0.1% Tween 20, 0.01% $NaN_3$) for 30 min at RT. The membrane was incubated for 1 hour with primary antibodies (see Table S3) at room temperature and then 1 hour with secondary antibodies conjugated with either IRDye 700DX or IRDye 800CW (LI-COR). The membrane was washed with PBS containing 0.1% Tween 20 after adding each antibody and was imaged using Odyssey CLx (LI-COR).

For qPCR analysis, total RNA was isolated using RNeasy kit (Qiagen) and treated with DNase I (DNA free kit, Ambion) to remove genomic DNA. The efficient removal of genomic DNA in the RNA samples was tested by performing PCR using primers against the *GAPDH* genomic locus. Synthesis of cDNA was performed using qScript cDNA Supermix (Quanta Biosciences) from 1 µg of total RNA according to the manufacturer's protocol. cDNA was diluted 100x and 2.5 µl aliquots subjected to qPCR with SensiMix SYBR & Fluorescein Mastermix (Bioline) and appropriate primers (see Table S4) on a LightCycler 480 (Roche).

For immunofluorescence analysis, cells were seeded on coverslips and either fixed the next day for undifferentiated LUHMES cells or differentiated for defined time points and then fixed with 4% formaldehyde for 10 min at room temperature. Cells were permeabilised with 0.2% Triton X in PBS for 10 min and blocked with 10% fetal bovine serum in PBS for 30 min at RT. Primary antibody (see Table S3) incubation was performed in 1% FBS, 0.1% Tween 20 in PBS for 1 hour at RT and coverslips were washed three times with 0.1% Tween 20 in PBS. Secondary antibodies (Alexa Fluor 488, Alexa Fluor 555, Alexa Fluor 647, Thermo Scientific) were diluted 1000x in 1% FBS, 0.1% Tween 20 in PBS and applied for 1 hour at RT. Finally, cells were washed three times with 0.1% Tween 20 in PBS, stained with 5000-fold diluted DAPI in PBS for 10 min at RT, mounted using Prolong Gold or Diamond (Thermo Scientific) and dried overnight at RT. Cells were imaged on the Leica TCS Sp5 confocal microscope (Leica Microsystems) using either 40x or 63x oil immersion objectives. Image analysis was performed using Volocity (PerkinElmer). Live cell imaging of differentiating neurons was performed using the IncuCyte Zoom system (Essen BioScience) and neurite lengths were analysed using NeuroTrack package from the IncuCyte software.

##### 1.4. Preparation of total DNA for HPLC.

Cells were washed in PBS, lysed in lysis buffer (10mM Tris HCl [pH 7.4], 50 mM NaCl, 0.5% SDS, 100mM EDTA, 300 µg/ml proteinase K) and incubated at 50°C for 2 hours. Total nucleic acid was recovered from the completely lysed sample by ethanol precipitation in 2 volumes of 100% ethanol at room temperature (for 30 minutes), and pelleting by centrifugation. The pellet was washed once in 2 volumes of 70% ethanol, and the nucleic acid pellet was resuspended in hydrolysis buffer containing 1x DNase I buffer (*NEB*), 1mM zinc sulphate, DNase I (*NEB*) and Nuclease P1 (*Sigma*). After 4 hours, the sample was mixed thoroughly and digested for a further 8 hours. After 12 hours at 37°C, the sample was heated to 92°C for 3 minutes and cooled on ice. Two volumes of 30mM sodium acetate, 1mM zinc sulphate [pH 5.2] were added plus additional Nuclease P1 and the nucleic acids were further digested to deoxyribonucleotide and ribonucleotide 5' monophosphates for a further 24 hours at 37°C. The samples were then subjected to HPLC.

##### 1.5. HPLC quantification of nucleotide content.

HPLC was performed on the 5 $\mu$m Apex ODS C18 column, with isocratic 50 mM ammonium phospate (monobasic) mobile phase. UV absorbance was recorded at 276 nm (dCMP, elution time 9.4 minutes), 282 nm (5mdCMP, elution time 17 minutes), 268 nm (dTMP, elution time 21.9 minutes), 260nm AMP and dAMP (elution times 27 minutes and 62.47 minutes) and 254 nm (GMP and dGMP, elution times 11.1 minutes and 29.7 minutes). Extinction coefficients used in nucleotide quantifications were dCMP, 8.86 x $10^3$; 5mdCMP 9.0 x $10^3$; dTMP, dGMP/GMP 12.16 x $10^3$; dAMP/AMP 15.04 x $10^3$. Relative amounts of all nucleotides were calculated from the area under each peak (Chromeleon software) using the respective extinction coefficients.

##### 1.6 Methylation-dependent repression assay.

Repression assay was performed using Dual Luciferase assay kit (Promega) according to manufacturer's protocol. First, we inserted Firefly luciferase containing CpGs into CpG-free plasmid obtained from InvivoGen. 100 µg of CpG-free luciferase plasmid was methylated with 200 U M.Sss I (NEB) in the presence of SAM for 2 h 40 min at 37°C. Simultaneously, the same amount of the plasmid was incubated in the buffer with SAM but without M.Sss I to be used as unmethylated control. Reaction was deactivated at 65°C for 20 min. Plasmids were purified from proteins by PCI (Sigma) and DNA was precipitated from the water phase using isopropanol. To confirm the methylation status of the plasmids, restriction enzymes Hpa II and Msp I (NEB) were used. CpG-free luciferase plasmid, plasmids containing human MeCP2 either WT or R111G or R306C and plasmid containing *Renilla* luciferase were transfected into *MBD2-/- MeCP2 -/y* MEFs (total amount transfected: 500 ng). Specifically, 5 ng of unmethylated or methylated CpG-free luciferase plasmid was mixed with 500 ng of CpG- and luciferase-free plasmid, and further mixed with 100 ng of plasmid expressing MeCP2 and 12.5 ng of plasmid expressing *Renilla* luciferase. This mixture of DNA was combined with 3.5 µl of Lipofectamine 2000 (Thermo Scientific) in the OPTIMEM medium and added onto MEFs seeded on the day before transfection at the density of 50,000 cells per well of a 24-well plate. The transfection mixture was incubated with cells at 37°C for 5-6 hours and after that the medium was changed. Dual luciferase assay was performed 48h after transfection as follows. Transfected cells were lysed in 1x Passive Lysis buffer at RT for 15 min with gentle rocking. 100 µl of Luciferase substrate were transferred into an illuminometer tube. 20 µl of lysed cells were added and luminescence of firefly luciferase was recorded. Next, immediately after the first measurement of firefly luciferase, Stop and Glo reagent was added and *Renilla* luciferase activity was measured. All measurements were done in at least three replicates and ratios of Firefly luciferase and *Renilla* luciferase activities were calculated for unmethylated and methylated Firefly luciferase.

#### 2. Library preparation for Next Generation Sequencing.

##### 2.1. TAB-seq.

The TAB-treated genomic DNA was sonicated for 30 cycles of 30 sec ON and 30 sec OFF on low power using Bioruptor (Diagenode). DNA was end-repaired and the ends were 3'-adenylated in order to facilitate adapter ligation. Size selection was performed using Agencourt AMPure XP (Beckman Coulter) beads. After adapter ligation and size selection, DNA was treated using the EpiTect Bisulfite kit (Qiagen) and PCR amplified using custom primers. All libraries were sequenced as 100 bp long pair-end reads on HiSeq 2500 Illumina platforms.

##### 2.2. RNA-seq.

Total RNA was isolated from all generated cell lines (Table S1) at day 9 of differentiation using either the RNeasy Mini kit or the AllPrep DNA/RNA Mini kit (Qiagen). Genomic DNA contamination was removed with the DNA-free kit (Ambion) and remaining DNA-free RNA was tested for purity using PCR for the *GAPDH* genomic locus. Total RNA was tested on the 2100 Bioanalyzer (Agilent Technologies) to ensure a RIN quality higher than 9, and quantified using Nanodrop. Equal amounts of total RNA were taken forward for library preparation and ERCC RNA Spike-in control mixes (Ambion) were added according to the manufacturer's guide. Ribosomal RNA was depleted using the Ribo-Zero Gold rRNA Removal module (Epicentre, Illumina). Isolated mRNA was tested for purity on the 2100 Bioanalyzer. mRNA was quantified using Qubit and equal amounts of each sample were used for cDNA synthesis and 3' terminal tagging using ScriptSeq v2 RNA-seq library preparation kit (Epicentre, Illumina). Libraries were PCR amplified to add adaptors and barcodes. The libraries were sequenced as 100 bp pair-end reads using HiSeq 2000 or HiSeq 2500 Illumina platforms.

##### 2.3. ATAC-seq.

Neurons were scraped from the plate and nuclei were isolated using a hypotonic buffer (10 mM Tris-HCl pH7.4, 10 mM NaCl, 3mM $MgCl_2$, 0.1% [v/v] Igepal CA-630), and counted. 50,000 nuclei were resuspended in 50 µl of a transposition reaction mix containing 2.5 µl Nextera Tn5 Transposase and 2x TD Nextera reaction buffer. The mix was incubated for 30 min at 37 ºC. DNA was purified by either the MinElute PCR kit (Qiagen) or the Agencourt AMPure XP (Beckman Coulter) beads and PCR amplified with the NEBNext High Fidelity reaction mix (NEB) to generate DNA libraries. The libraries were sequenced as 75bp long pair-end reads on a HiSeq 2500 Illumina platform.

##### 2.4. MeCP2 ChIP-seq.

LUHMES-derived neurons at day 9 of differentiation with four levels of MeCP2: KO, WT, OE 4x and OE 11x (Table S1) were crosslinked with 1% of Formaldehyde (Sigma) for 10 min at room temperature (RT) and quenched with 2.5 M Glycine (Sigma) for 2 min at RT. Cells were washed with PBS, scraped from the plate and centrifuged. Crosslinked nuclei were isolated in a hypotonic buffer (10 mM Tris-HCl pH7.4, 10 mM NaCl, 3mM $MgCl_2$, 0.1% [v/v] Igepal CA-630) and counted using a haemocytometer. Chromatin from ~$4x10^6$ nuclei was sonicated for 20 cycles (30 sec ON and 30 sec OFF) using Bioruptor (Diagenode) on high power. Crosslinked and sonicated chromatin was mixed with 60 ng of sonicated *Drosophila* chromatin (Active Motif) as a spike-in, and the mix was incubated overnight at 4 ºC with antibodies against MeCP2 (D4F3, Cell Signalling) plus spike-in antibody (Active Motif). After overnight incubation, magnetic Protein G coated beads (Thermo Scientific) were added and incubated for 4 hours at 4 ºC. Beads were washed, and chromatin was reverse-crosslinked overnight at 65 ºC. DNA was purified using the Agencourt AMPure XP (Beckman Coulter) beads. For ChIP-seq library preparation, IPs for each condition were pooled together to achieve 5 ng total DNA as a starting material. For example, 3-4 IPs were pooled together for the KO sample and 2 IPs were pooled for the WT sample. Libraries were prepared using the NEBNext Ultra II DNA library Prep kit (NEB) for both IPs and corresponding inputs. The libraries were sequenced as 75bp long pair-end reads on a HiSeq 2500 Illumina platform.

#### 3. Bioinformatics and data preparation.

##### 3.1. Bisulfite sequencing (TAB-seq).

Trimmomatic version 0.32 (33) was used to perform quality control on 94b and 75 bp paired-end reads to remove adapter sequence and poor quality bases at the ends of reads for both BS-sq and ChIP-seq. For BS-seq, we used Bismark version 0.10 (34) to further align and process the reads. Mapping was performed in bowtie2 mode to the human hg19 reference genome. Following alignment, reads were deduplicated and methylation values were extracted as bismark coverage and cytosine context files. We calculated the methylation percentage at each cytosine position as (mC/C)x100 and generated *.bed files for further processing.

##### 3.2. ChIP-seq and ATAC-seq.

We used bwa mem version 0.7.5 (38) to map reads to the human hg19 reference genome. We filtered the alignments to remove reads that map to multiple locations in the genome and to blacklisted regions defined by the ENCODE project. We further removed duplicate reads with Picard version 1.107 MarkDuplicates (http://broadinstitute.github.io/picard/). To account for varying read depths we used deepTools version 2.5.1 (39) to create bigWig files normalised by RPKM (reads per kilobase per million reads). To quantify MeCP2 occupancy on the genomic features of interest (mCG, mCA, GT, etc.), we reject reads longer than 1 kb regarding them as alignment artefacts.

##### 3.3. RNA-seq.

All paired-end sequencing reads were trimmed, and quality controlled using Trimmomatic version 0.33 (33). The filtered reads were then mapped using STAR version 2.4.2 (40) using hg19 human genome assembly and Ensembl 74 release for annotation. Additionally, TPM values for genomic features were quantified by quasi-mapping approach using Sailfish version 0.10.0 (41). Protein coding transcripts for Sailfish index generation were taken from Gencode release 19. In order to assign reads to genomic features, featureCounts version 1.5.0 was used (42). Differential Expression analysis was performed using DESeq (43). Comparisons between mutant vs wildtype and were performed within the same batch using the design formula ~differentiation+condition in order to account variance from the differentiation.

#### 4. ChIP-seq accumulation algorithm.

For each chromosome we create an array $n_i$ of the number of instances a particular locus (a single base pair at position $i$ in the chromosome) is covered by a ChIP-seq read. We do this by going through all ChIP-seq reads from *.bed files, each time incrementing all $n_i$'s for which $i$ is between the start and end position of a given read. When all reads in a given chromosome have been processed, for each feature $x$ ($x$ =mCG, mCA, …) at position $j$ we select a region of interest of length $L$ around it ($j \pm L/2$ bp) and accumulate the counts in a separate array (different for each feature $x$): $c_i^x \to c_i^x + n_{i+j}$ , for all $i$ such that $-\frac{L}{2} \leq i < \frac{L}{2}$. The obtained accumulated counts $c_i^x$ for ChIP-seq are divided by the accumulated counts for the corresponding input DNA-seq to reduce the sequencing bias present in the data.

To demonstrate the above procedure, let us take the following sequence as the reference genome with features of interest (in this case CG) marked red:

```
aaacagctagttaattttgaatcgcaggtaaacaatcgaataattttcta
```

We assume that ChIP-seq has generated the following reads:

```
 ttttgaatcgca
      aatcgcaggta
attttgaatcg
         cgcaggtaa
                   caatcgaa
                      tcgaataattt
```

The number of reads $n_i$ covering a specific site in the genome is

```
0000000000000122222333443322221011122221111110000
```

Raw counts over a fixed-length region (here +/-5bp, in the actual algorithm this would be +/-5000bp) centred at the feature of interest are

```
22333443322 2
011122222111
```

Total accumulated counts $c_i^{CG}$ versus position $i$ relative to the feature of interest is therefore

```
23445665533 3
```

In the actual analysis, the above sequence would be a 10kbp-long array of integers.

#### 5. Details of the ChiP-seq computer model.

##### 5.1. The algorithm.

The input to the simulation are the *.bed files with ChIP or input (DNA) reads, and the two parameters $p, p_{bg}$. The simulation uses the data files to preserve the distribution of the lengths of reads but not their positions. The positions are determined by the following algorithm:

1) For each input file, GC content- and length bias is first estimated by constructing a table of counts $r[\%GC][l]$ for the actual reads and $g[\%GC][l]$ for genomic sequences of the same length $l$, %GC, and randomly selected locations.
2) For each read from the experimental *.bed file we calculate its length $l = e - s$ and then select a random start point $s$ (within the same chromosome as the original read) and the end point $e = s + l$.
3) GC content and the number of mCs for the simulated read is calculated. We take a particular C to be methylated with probability proportional to the fraction of methylated reads containing this site in our methylation data (TAB-seq).
4) If there is at least one mC within the read, an auxiliary variable $P$ is set to 1 with probability $p$ times the relative binding affinity of the motif to which this C belongs. Otherwise $P$ is set to $p_{bg}$. This accounts for MeCP2 present/absent in this particular DNA fragment.

5) $P$ is multiplied by $r[\%GC][l]/g[\%GC][l]$ to account for the GC content and read length bias.
6) The simulated read is accepted with probability $P$ and saved to a file, or rejected (with probability $1-P$). If the read is rejected, another random position $s$ is selected and the algorithm continues from (3).
7) Go to step 2 unless all reads from the *.bed file have been processed.

Simulated reads are then processed in the same way as the experimental ChIP data (SI Section 4).

##### 5.2. Height of the ChIP-seq enrichment peak.

Since mCG is much more frequent than any other MeCP2-binding motif in our cell lines, in what follows we focus on enrichment profiles on mCG. Figure SI 1, left, shows simulated enrichment profiles on mCG obtained for $p_{bg}=0$ and different $p$. Counterintuitively, the height of the central peak does not decrease with decreasing $p$, but it slowly increases. The height of the peak tends to a constant as $p \to 0$ (Figure SI 1, right). This can be explained as follows. For large $p$, fragments centred at adjacent mCGs overlap, increasing the counts $c_i^{mCG}$ in the flanking regions ($|i| \gg 1$). Normalization lowers the apparent height of the peak since it divides the profile by the counts in the flanking regions. As $p$ decreases, the number of overlapping fragments from neighbouring mCGs decreases; this reduces raw counts in the flanks and increases the relative height of the central peak after normalization.

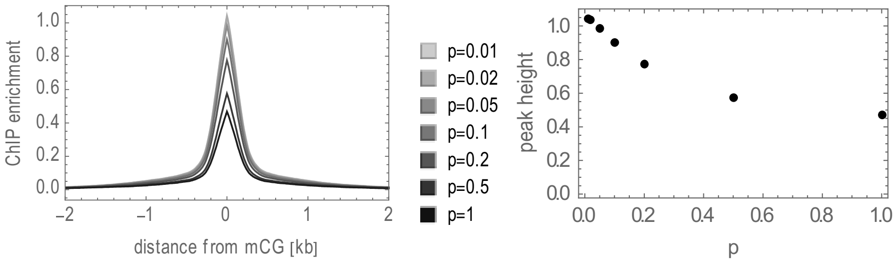


Figure SI 1 Simulated ChIP enrichment profiles for different DNA-MeCP2 attachment probabilities $p$ show a counter-intuitive reciprocal dependence of profile's height on MeCP2 occupancy.

##### 5.3. Fitting ChIP-seq to data.

For each ChIP-seq data set we fitted the simulated profile $f_i^{sim}$ (parametrized by $p, p_{bg}$) to the experimental enrichment profile $f_i^{expr}$ by minimizing the sum of squared differences,

$$\chi^2 = \sum_{i=-100}^{100} \left(f_i^{expr} - f_i^{sim}\right)^2,$$

with respect to $p$ and $p_{bg}$. The region +/-100bp was chosen because of the sensitivity of the profile shape to $p, p_{bg}$ in this region. Minimization was performed as follows: we simulated profiles for a range of $p$ and two values of $p_{bg}$: 0 and 0.9; profiles for $p_{bg}$ between these two values were obtained by linear interpolation (permitted due to the assumed additive model of signal and background reads). The minimum $\chi^2$ obtained in this way for 11x OE is plotted in Fig. S5D. Any $p \leq 0.1$ gives similar (low) values of $\chi^2$, indicating that $p \approx 0.1$ is the upper bound on mCG occupancy in 11x OE.

To predict profiles on features other than mCG (Figs. S5E,F), we used the values of $(p, p_{bg})$ obtained from the mCG profiles by minimizing $\chi^2$ with respect to $p_{bg}$ and assuming $p = 0.1, 0.05, 0.02, 0.002$ for 11x OE, 4x OE, WT, and KO, respectively. These values are based on the best-fit $p = 0.1$ obtained for 11x OE, the other $p$'s being a fraction of this value approximately proportional to the relative abundance of MeCP2 in a given cell line versus 11x OE. The predicted profiles match the data very well but are not sensitive to the exact value of $p$. This indicates that ChIP-seq alone cannot be used to estimate the occupancy $p$.

### 6. ATAC-seq model.

#### 6.1. The algorithm.

We simulate binding to a short DNA sequence (first $L$=50Mbp of chromosome 1). We assume that MeCP2 occupies 11bp (20, 21) and that the protein is centred on an mC, thus the obscured genomic sequence is xxxxxmCxxxxx where x can be any nucleotide.

The following algorithm is repeated $T$ times (larger $T$ corresponds to longer digestion times in the actual experiment):

1) A random location $i$ is chosen as the position of the new cut. We assume the following convention: $i$ refers to the position of the nucleotide immediately to the left from the centre of the cut. Hence, $i+1$ denotes the position of the nucleotide to the right from the cut. Positions $i, i+1$ are the positions of insertion sites.
2) The position is accepted with probability $p_i$ which depends on the nucleotide sequence $i-w, \ldots, i+w$ where $w$=10 bp. This is based on the known Tn5 sequence preference across $21 = 2w+1$ bp that it contacts (19, 44). The probability $p_i$ is calculated using the position weight matrix (PWM) obtained from ATAC-seq reads for KO1.

3) If the position is rejected, go back to 1.
4) If there is enough space for Tn5, a cut is made between $i$ and $i + 1$. We assume that Tn5 occupies 21bp $(i - w, \dots, i + w)$ and for a cut to be made there must not be any protein (MeCP2) overlapping with this region. There must not be any previous cuts made in this region too.
5) If the proposed cut is rejected, go to step 1 and try again.
6) A weight $W = \exp(b \times GC)$ is assigned to each fragment where $GC$ is the GC content (0…1) of the fragment and $b$ is a constant. This simulates an additional GC bias that may be created by the sequencing procedure.

The shortest possible fragment generated by the algorithm is 21bp which corresponds to two Tn5 cutting just next to each other. Since MeCP2 occupies 11bp, the shortest MeCP2-containing fragment is 11+21=32bp. We retain only fragments longer than 35bp because our experimental data does not contain shorter fragments.

The simulation creates artificial fragments and their weights that we process in the same way as the experimental data. The input to the program is the DNA sequence (fixed), the density $p$ of MeCP2 on mCxx, the average density of insertion (cut) sites $t = T/L$ (cannot be larger than 1/21bp), and the GC bias $b$.

##### 6.2. Algorithm benchmarking

To test the role of the parameters on the shape and depth of the footprint of MeCP2, we simulated the model for different $p, t, b$. We also performed simulations with/without Tn5 bias.

Figure SI 2a shows the counts profiles for $p = 0$ (no MeCP2) with and without Tn5 bias, compared to experimental profiles. It is evident that the insertion bias is required to reproduce the experimental insertion profiles. However, the bias cancels out when calculating the relative profile (footprint) $f_i$. This is demonstrated in Figure SI 2b which shows the footprint obtained by dividing the counts profile for $p = 0.05$ by the profile for $p = 0$, for different $t$ and $b = 6$ (Tn5 bias) or $b = 0$ (no bias). Increasing digestion time $t$ increases the height of the side peaks surrounding a depression caused by MeCP2. However, the depth of the depression does not change noticeably.

Figure SI 2c, left shows that changing the GC bias $b$ can does not change the depth of the footprint if the bias is the same in the simulated test and reference samples. However, the footprint can change if the GC bias is different in the two samples, see Figure SI 2c, right. Since such a mismatch in the GC bias would have a notable signature (rising/falling flanks

>50bp away from mCG) which we do not see in our data, we conclude that the bias is very similar in all experimental samples.

Figure SI 2d shows that dividing the counts for $p > 0$ by the counts for $p = 0$ but with a different digestion time $t$ changes the depth of the footprint slightly. Experimental variation causes $t$ to be slightly different for different samples, and hence our results can have a small systematic error.

We also simulated the presence of nucleosomes by treating them as another protein of size 147bp that binds to the DNA, and distributing them randomly and uniformly on the DNA. We found no differences in the footprint compared to the case with no nucleosomes (data not shown). However, the distribution of the number of fragments exhibited maxima at fragment lengths equal to runs of one, two, three etc. nucleosomes. This is in qualitative agreement with the ATAC-seq fragment length distribution (data not shown).

Our simulations cannot explain two subtle features of the experimental data: the presence of oscillations superimposed on the main profile, and a small central peak visible in WT/KO, CTRL/KO, and OE 4x/KO. We speculate that the first is caused by steric interactions between MeCP2 and Tn5, and the latter by interactions of proteins other than MeCP2 with mC or a small difference in the GC bias among the samples.

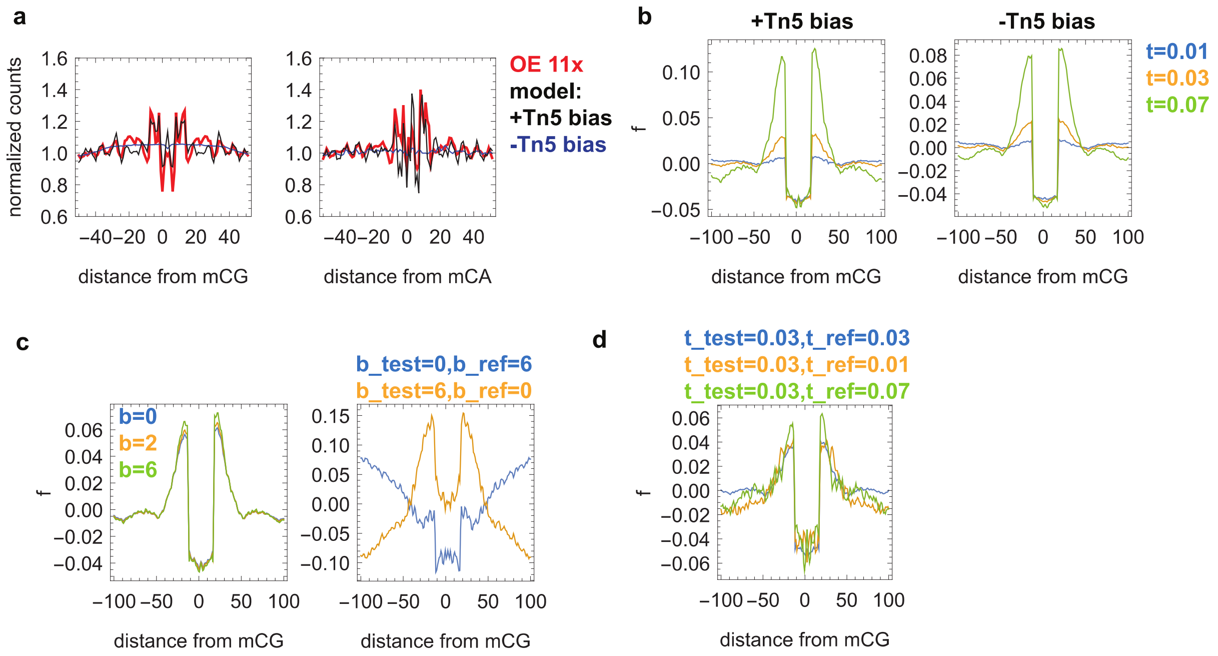


Figure SI 2 Benchmarking of the simulated ATAC-seq. (a) Comparison between experimental and simulated $(p = 0, t = 0.04, g = 6)$ ATAC-seq counts shows that Tn5 bias is important in reproducing the normalized counts *in silico*. Red = OE 11x, blue = simulated with no Tn5 bias, black = simulated with Tn5 bias. (b) Simulated footprint $f$ for $p = 0.05, g = 6$ and $t =$

$0.01, 0.03, 0.07$ (blue, yellow, and green respectively) shows that excluding the Tn5 insertion bias does not significantly affect the depth of the footprint. Left = with Tn5 bias, right = no bias. (c) The role of CG bias $b$ on the simulated footprint $f$. CG bias cancels out if identical in both test and reference samples. Left: the same $b = 0,2,6$ (blue, yellow, and green respectively) for the test and reference samples. The footprint is not affected by the bias. Right: different $b$'s for the test and reference samples: $b_{test} = 0, b_{ref} = 6$ (blue) and $b_{test} = 6, b_{ref} = 0$ (yellow). The shape and depth of the footprint is significantly affected. In all cases $p = 0.04$. (d) The shape of the footprint depends on the difference in digestion time $t$ between the test and the reference sample, but the depth of the footprint does not. Blue = $(t_{test} = 0.03, t_{ref} = 0.03)$, yellow = $(t_{test} = 0.03, t_{ref} = 0.01)$, green = $(t_{test} = 0.03, t_{ref} = 0.07)$. In all cases, $p = 0.05, b = 6.0$.

##### 6.3. Fitting simulated ATAC-seq profiles to data.

We used the depth of the footprint to extract MeCP2 occupancy $p$. We simulated ATAC-seq for many pairs $(p, t)$, for $p = 0 \dots 0.1$, $t = 0 \dots 0.08$, and a fixed $b = 6.0$ (the exact value is not important since GC bias cancels out when calculating $f_i$). We then selected those $(p, t)$ which minimized the distance between the simulated and experimental footprints $f_i^{sim}$ and $f_i^{exp}$,

$$D = \sum_{i=\{-9,-10,14,15\}} \left(f_i^{exp} - f_i^{sim}\right)^2 \times \sum_{i=-100}^{100} \left(f_i^{exp} - f_i^{sim}\right)^2.$$

This formula assigns a larger weight to the region (edges of the central dip) which is least sensitive to variations in the ATAC-seq protocol, and reduces the systematic error caused by the (small) central peak of unknown origin (SI Section 6.2). The best fit to the 11x OE footprint yields $p = 0.063, t = 0.04$. Fig. 2E shows $p$ for all MeCP2 levels $M$. The relationship is linear, with the best-fit $p = 0.0058 \times M_{cell\ line}/M_{WT}$.

#### 7. Generic model of gene expression.

Below we list all the parameters and observables of the generic ("paradigm") model of gene expression (22) from Fig. S7A which is the basis for all other models. For each gene $i$ we define

- $k_{i,ON}, k_{i,OFF}$ – the switching rates OFF→ON and ON→OFF, respectively.
- $J_i$ – transcription rate (in transcripts/second).
- $\alpha_i$ – transcription initiation rate (1/second)

- $v_i$ - transcription elongation rate (bp/second).
- $d_i$ – mRNA degradation rate (1/second).
- $f_i$ – the fraction of cells in the population that at any given time have gene $i$ in the ON state.

We assume that mRNA is degraded according to 1st order kinetics (45). The steady-state concentration $c_i$ of mRNA from gene $i$ is obtained by equating production (transcription in the ON state) and degradation rates:

$$J_i f_i = c_i d_i, \qquad \text{(Equation S1)}$$

from which it follows that

$$c_i = J_i f_i \,/d_i. \qquad \text{(Equation S2)}$$

The number of transcripts per million bp (TPM) which we obtain from RNA-seq is thus

$$\mathrm{TPM}_i = \; N J_i f_i \,/d_i, \qquad \text{(Equation S3)}$$

where $N$ is an unknown proportionality constant that depends on the sample and is different for different cell lines, replicates and conditions but is the same for all genes in a given sample.

#### 8. Condensation model.

This model considers a hypothesis (ultimately proven to be false, see the main text) that MeCP2 causes chromatin condensation which reduces the fraction of cells with genes in the active (ON) state (Fig. 3A). We assume that the fraction $f_i$ of cells with gene $i$ in the active state depends on promoter openness $a_i$ (measured by ATAC-seq) which in turn depends on the level $M$ of MeCP2 and gene methylation $\rho_i$:

$$f_i = f_i(M, \rho_i) \propto a_i = a_i(M, \rho_i). \qquad \text{(Equation S4)}$$

The model also assumes that transcription rate $J_i$ and mRNA degradation rate $d_i$ are not affected by MeCP2. The Log2FC of differential expression of gene $i$ for cell lines X and Y is then

$$\mathrm{Log2FC}_{\mathrm{X/Y},i} = \log_2 \frac{\mathrm{TPM}_i(\mathrm{X})}{\mathrm{TPM}_i(\mathrm{Y})} = \log_2 \frac{N_\mathrm{X}}{N_\mathrm{Y}} + \log_2 \frac{a_i(M_\mathrm{X}, \rho_i)}{a_i(M_\mathrm{Y}, \rho_i)}.$$

The average Log2FC of genes with the same methylation density $\rho$ is therefore

$$\mathrm{Log2FC}_{\mathrm{X/Y}}(\rho) = \log_2 \frac{N_\mathrm{X}}{N_\mathrm{Y}} + \langle \log_2 \frac{a_i(M_\mathrm{X}, \rho)}{a_i(M_\mathrm{Y}, \rho)} \rangle,$$

where ⟨…⟩ denotes averaging over genes with the same $\rho$. According to the above equation, $\mathrm{Log2FC}_{\mathrm{X/Y}}$ should yield the same curve (modulo a vertical shift due to different normalization factors $N_\mathrm{X}$ and $N_\mathrm{Y}$) as the logarithm of the ratio of accessibilities of X versus Y when plotted as a function of methylation density. Figure 3C shows that this is not the case. We can therefore reject the hypothesis that MeCP2 modulates gene expression primarily by altering the fraction of active genes.

#### 9. Detachment model.

In this hypothetical scenario we assume that RNA Pol II aborts transcription with some small probability $\lambda$ when it collides with MeCP2 or encounters a chemical mark left by the interaction between MeCP2 and chromatin. The probability that RNA Pol II reaches the end of the gene (transcription end site, TES) is thus

$$P = (1-\lambda)^n \cong e^{-\lambda n},$$

where $n$ is the number of "abort sites" on the gene, proportional to the number of MeCP2 molecules on the gene. ATAC-seq shows that the density of MeCP2 on a gene is proportional to its methylation density $\rho$ and the total concentration $M$ of MeCP2 in the nucleus, hence we can write that $n = AM\rho L = AMN_{\mathrm{mCG}}$, where $A$ is an unknown proportionality factor, $L$ is the length of the gene, and $N_{\mathrm{mCG}}$ is the total number of mCGs. Log2FC of the differential expression X versus Y can then be written as

$$\mathrm{Log2FC}_{\mathrm{X/Y}} = \log_2 \frac{\mathrm{TPM}_i(\mathrm{X})}{\mathrm{TPM}_i(\mathrm{Y})} = \log_2 \frac{N_\mathrm{X}}{N_\mathrm{Y}} + \log_2 \frac{P(M_\mathrm{X}, N_{\mathrm{mCG}})}{P(M_\mathrm{Y}, N_{\mathrm{mCG}})}$$

$$= \mathrm{const} + \log_2 \frac{\exp(-\lambda A M_\mathrm{X} N_{\mathrm{mCG}})}{\exp(-\lambda A M_\mathrm{Y} N_{\mathrm{mCG}})}$$

$$= \mathrm{const} - \gamma \ \Delta M_{\mathrm{X/Y}} N_{\mathrm{mCG}},$$

where $\gamma = \lambda A M_\mathrm{Y} / \log_2 e$ is an unknown parameter identical for all cell lines, and $\Delta M_{\mathrm{X/Y}} = \frac{M_\mathrm{X}}{M_\mathrm{Y}} - 1$ is the relative difference between the level of MeCP2 in cell lines X and Y. For example, $\Delta M_{\mathrm{KO/WT}} = -1$ and $\Delta M_{\mathrm{11x\,OE/WT}} = 10$. $\mathrm{Log2FC}_{\mathrm{X/Y}}$ should therefore follow a straight line when plotted versus $N_{\mathrm{mCG}}$, and the slope of this line should be positive for KO/WT, and negative (and 10x more steep) for 11xOE/WT. Figure 3E shows that when the model is fitted to the KO data to fix the unknown constant $\gamma$, it fails to reproduce the OE 11x data. We hence conclude that there is no evidence that MeCP2 cause premature termination of transcription.

### 10. Congestion models.

#### 10.1. General considerations.

We consider a hypothesis that MeCP2 slows down the elongation step of transcription by creating queues of RNA Pol II in front of MeCP2 or chemical modifications left by it in gene bodies (Figs. 4A and S7B). We assume that the density of obstacles, or "slow sites", is proportional to the density of molecules of MeCP2 bound to the DNA.

We show in the main text that (i) the density $m$ of MeCP2 on the DNA is proportional to the concentration $M$ of MeCP2 in the cell and (ii) that $m$ is proportional to mCG density $\rho$. We can thus write that the transcription rate $J$ averaged over many genes with the same mCG density $\rho$ is

$$J = J(M\rho, \rho).$$

The second argument of $J$ accounts for non-MeCP2 but mCG-density dependent modulation. Figure SI 3 show that gene expression (TPMs) versus gene body methylation for KO and 11x OE. The difference between the two cells lines is very small, and non-MeCP2 dependent component of $J$ dominates the behaviour of $J(\rho)$. We are thus permitted to rewrite $J$ in the factorized form

$$J \approx [1 - \epsilon(M\rho)]K(\rho),$$

where $K$ is a fast-changing function of $\rho$ and the correction $\epsilon(m)$ due to MeCP2 is a slowly increasing function of $m$, and $\epsilon(0) = 0$. This leads to the following expression for the Log2FC of cell lines X versus Y:

$$\mathrm{Log2FC}_{\mathrm{X/Y}}(\rho) = \log_2 \frac{\mathrm{TPM}_i(\mathrm{X})}{\mathrm{TPM}_i(\mathrm{Y})} \approx \log_2 \frac{N_{\mathrm{X}}}{N_{\mathrm{Y}}} + \log_2 \frac{[1 - \epsilon(M_{\mathrm{X}}\, \rho)]K(\rho)}{[1 - \epsilon(M_{\mathrm{Y}}\rho)]K(\rho)}$$

$$= \mathrm{const}(\mathrm{X}, \mathrm{Y}) + \log_2[1 - \epsilon(\rho M_{\mathrm{X}})] - \log_2[1 - \epsilon(\rho M_{\mathrm{Y}})$$

$$\approx \mathrm{const}(\mathrm{X}, \mathrm{Y}) - \epsilon(\rho M_{\mathrm{X}}) + \epsilon(\rho M_{\mathrm{Y}}).$$

The latter approximation is valid since $\epsilon$ is supposed to be small. In particular, the Log2FC of any cell line X versus KO reads

$$\mathrm{Log2FC}_{\mathrm{X/KO}}(\rho) \approx \mathrm{const}(\mathrm{X}, \mathrm{KO}) - \epsilon(\rho M_{\mathrm{X}})$$

because $\epsilon(\rho M_{\mathrm{KO}}) = \epsilon(0) = 0$. The Log2FC curves for different cell lines versus KO will therefore have the same shape when plotted in the variable $\rho M_{\mathrm{X}}$. It follows from this that the maximum slope of these lines will be proportional to $M_{\mathrm{X}}$.

A similar argument holds when the reference is not the KO but a control line CTR with low but non-zero level of MeCP2, following a mild assumption that $\epsilon(m)$ is linear in $m$ for small $m$ and saturates for large $m$. We have

$$\text{Log2FC}_{\text{CTR/KO}}(\rho) \approx \text{const}(\text{CTR, KO}) - \epsilon(\rho M_{\text{CTR}}).$$

We can calculate $\epsilon(m)$ from this equation:

$$\epsilon(m) \approx \text{const}(\text{CTR, KO}) - \text{Log2FC}_{\text{CTR/KO}}(m/M_{\text{CTR}})$$

This enables us to express the Log2FC of any cell line X versus control CTR as

$$\text{Log2FC}_{\text{X/CTR}}(\rho) \approx \text{const}(\text{X, CTR}) - \epsilon(\rho M_{\text{X}}) + \epsilon(\rho M_{\text{CTR}}).$$

$$= \text{const}(\text{X, CTR}) + \text{Log2FC}_{\text{CTR/KO}}(\rho M_{\text{X}}/M_{\text{CTR}}) - \text{Log2FC}_{\text{CTR/KO}}(\rho)$$

The maximum slope will occur for $\rho = 0$. In the vicinity of this point,

$$\text{Log2FC}_{\text{X/CTR}}(\rho) \approx \text{const}(\text{X, CTR}) + \rho M_{CTR}\epsilon'(0)\left(\frac{M_{\text{X}}}{M_{\text{CTR}}} - 1\right),$$

where $\epsilon'(0)$ is the derivative of $\epsilon(m)$ at $m = 0$. The maximum slope is therefore proportional to $M_{\text{X}}/M_{\text{CTR}} - 1$. This is corroborated by experimental results presented in Fig. 1C,D.

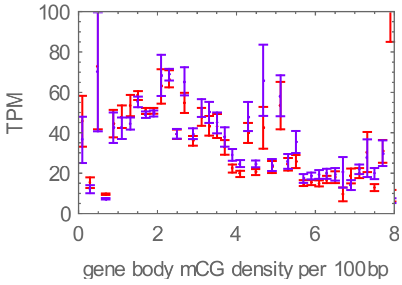


Figure SI 3 TPM (transcripts per million) as a function of gene body methylation density, for KO (violet) and OE 11x (red).

#### 10.2. Slow sites model.

In this model, MeCP2 causes a chromatin modification that slows down the RNA polymerase II. To mathematically model this process we use totally asymmetric simple exclusion process (TASEP) with open boundaries (24, 25, 46). A gene is represented as a one-dimensional chain of $L$ sites. Each site is either occupied by a particle, representing RNA Pol II, or is empty. Particles enter the chain at site $i = 1$ with rate $\alpha$ (transcription initiation rate), move along the chain and exit at site $i = L$ with rate $\beta$. Sites can be either "fast" or "slow". Slow sites represent mCGs affected by the interaction with MeCP2, whereas fast sites are all other sites (methylated or not). The speed of the particles is $v$ on fast sites and $v_s$ on slow sites. Slow sites are randomly and uniformly distributed, and their density is $\rho_s = \rho p$ where $\rho$ is the mCG density, and $p$ is the probability that an mCG is occupied by MeCP2 (as in the ChIP-seq and ATAC-seq models). $p$ is taken to be 0.063 for OE 11x, and proportionally smaller for other cell lines ($p = 0.0058 M_{\text{cell line}}/M_{\text{WT}}$).

Since RNA Pol II occupies about 60bp on the DNA and moves with the speed of about 60-70 bp/s (47), it is convenient to equate a single site of the model with a 60bp-long stretch of the actual DNA, and set the RNA Pol II speed to $v = 1$ sites per second on fast sites. We also assume $\beta = v = 1$ sec$^{-1}$ so that RNA Pol II is not blocked from exiting the chain at the end ($\alpha < \beta$ for all genes).

We simulated this model for different chain lengths $L = 166, 500, 1666, 5000$ corresponding to gene lengths between 10kb and 300kb, and a range of initiation rates $\alpha \in [0.001,1]$, densities of slow sites $\rho_s \in [1/64, 8]$ (mCG density between 0.026 and 13.3 per 100bp), and slow-site velocities $v_s = 0.01, 0.02, 0.05, 0.1, 0.2$ (all rates are in 1/sec). For each set of $(\alpha, \rho_s, v_s)$ we first let the model to reach steady state ("thermalization step"). We then measured the flow $J$ of particles (equivalent to the rate of transcription) through the chain. The flow strongly depends on $\rho_s$ and only weakly (logarithmically) on the length $L$ (Figure SI 4, left and middle). Therefore, in what follows we fix $L = 5000$ sites, which is equivalent to the gene length of 300 kb. The flow $J$ obtained from these simulations is presented in Fig. S7C as a function of $\alpha$, for different mCG densities $\rho$, and for $p = 0.063$ (OE 11x). The flow is approximately linear in $\alpha$ until some critical $\alpha_c$ which depends on $\rho_s$, and saturates at $J = J_{max}$ when $\alpha > \alpha_c$.

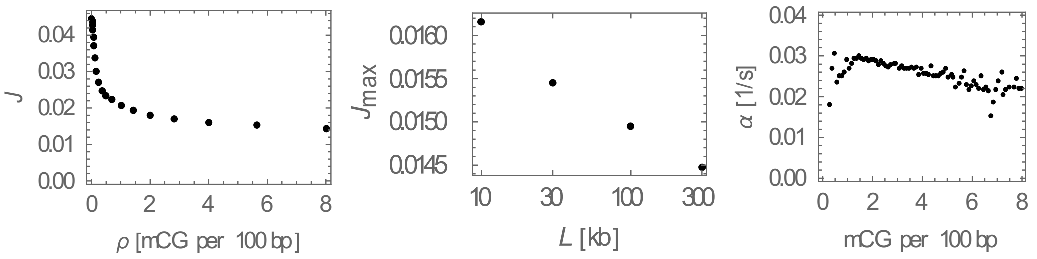


Figure SI 4 Left: Transcription rate $J$ as a function of mCG density, for $L = 30$ kb. Middle: maximum transcription rate ($\rho = 0$) versus gene length $L$. Right: estimated initiation rates as a function of mCG density.

To relate this model to the mRNA-seq differential expression data we must calculate Log2FC:

$$\text{Log2FC}_{\text{X/Y}} = \log_2 \frac{J(\alpha, \rho_{\text{s,X}})}{J(\alpha, \rho_{\text{s,Y}})},$$

where $\rho_{s,X} = \rho p_X$, $\rho_{s,Y} = \rho p_Y$ in which $\rho$ is mCG density and $p_X, p_Y$ are MeCP2 occupation probabilities for cell lines X,Y. We take $p_X = \left(\frac{M_X}{M_{OE11x}}\right) p_{OE11x} = 0.05 \left(\frac{M_X}{M_{OE11x}}\right)$, and similarly for $p_Y$. In the above expression we know all quantities except the initiation rate $\alpha$.

To obtain the initiation rate we bin genes according to their methylation as described, and then, for each bin, fit Log2FC(11xOE/WT) from the above equation to the experimental Log2FC. This gives us $\alpha(\rho)$ as a function of methylation density $\rho$ (Figure SI 4, right; average $\alpha = 0.027$ s$^{-1}$), as a calibration, to test our model prediction on the Log2FC data of the other cell lines. This approach, rather than fitting initiation rates of individual genes, removes correlations due to gene-gene interactions and produces a relatively smooth curve $\alpha(\rho)$. We can then use the fitted $\alpha(\rho)$ to predict $\mathrm{Log2FC}_{X/Y}$ for other pairs of cells lines. The results are presented in Figs. S7D,E and, as described in the main text, are in good agreement with experimental Log2FCs.

When the rates $\{\alpha\}$ are taken to be the same as for HeLa cells (1347 genes from (47) that were also present in our RNA-seq; average $\alpha = 0.05$ s$^{-1}$), the model still broadly agrees with the data, except for KO which is very sensitive to the exact values of $\alpha$ (plots not shown).

A model in which $J$ is first evaluated for individual genes (defined by their $\rho, L$) and then Log2FC obtained by binning genes according to their mCG density $\rho$ does not significantly affect the results (not shown).

##### 10.3. Dynamical obstacles model.

This model is very similar to the slow sites model with two exceptions: (i) polymerase always moves with the same speed $v$ (no slow sites) as long as it is not blocked by other polymerases and obstacles, (ii) obstacles binds and unbinds dynamically from the methylated sites. These obstacles can be MeCP2, other proteins recruited by MeCP2, or structural changes induced by MeCP2. We assume that unbinding occurs with rate $k_u$ per obstacle, whereas binding occurs with rate $k_u p$ per unoccupied mCG. An obstacle does not bind if an mCG is occupied by an obstacle or a polymerase. The parameters of the model are: $\alpha, v, L, \rho, p$ and, in addition, the unbinding constant $k_u$. Since the exact nature of obstacles in not specified, the density $p$ of sites that bind obstacles does not have to be the same as the MeCP2 occupancy estimated from ATAC-seq data. In fact, we found that the model reproduces the data best when $k_u = 0.04$, and $p = M/M_{OE11x}$, i.e., $p = 1$ for the OE 11x cell line.

The model behaves similarly to the slow-sites model. Fig. 4B shows a plot of the flow as a function of $\alpha$ and $\rho$. Fig. 4E,F shows that (after fitting the initiation rates as described for the slow-site model) the model is also able to reproduce the experimental data. The fact that the apparent fraction $p$ of occupied mCGs must be close to 1 in OE 11x suggest that MeCP2 may slow down RNA Pol II by altering chromatin structure rather than by direct steric interference.
